# Combination of a Sindbis-SARS-CoV-2 spike vaccine and αOX40 antibody elicits protective immunity against SARS-CoV-2 induced disease and potentiates long-term SARS-CoV-2-specific humoral and T-cell immunity

**DOI:** 10.1101/2021.05.28.446009

**Authors:** Antonella Scaglione, Silvana Opp, Alicia Hurtado, Ziyan Lin, Christine Pampeno, Maria G Noval, Sara A. Thannickal, Kenneth A. Stapleford, Daniel Meruelo

## Abstract

The COVID-19 pandemic caused by the coronavirus SARS-CoV-2 is a major global public threat. Currently, a worldwide effort has been mounted to generate billions of effective SARS-CoV-2 vaccine doses to immunize the world’s population at record speeds. However, there is still demand for alternative effective vaccines that rapidly confer long-term protection and rely upon cost-effective, easily scaled-up manufacturing. Here, we present a Sindbis alphavirus vector (SV), transiently expressing the SARS-CoV-2 spike protein (SV.Spike), combined with the OX40 immunostimulatory antibody (αOX40) as a novel, highly effective vaccine approach. We show that SV.Spike plus αOX40 elicits long-lasting neutralizing antibodies and a vigorous T-cell response in mice. Protein binding, immunohistochemical and cellular infection assays all show that vaccinated mice sera inhibits spike functions. Immunophenotyping, RNA Seq transcriptome profiles and metabolic analysis indicate a reprogramming of T-cells in vaccinated mice. Activated T-cells were found to mobilize to lung tissue. Most importantly, SV.Spike plus αOX40 provided robust immune protection against infection with authentic coronavirus in transgenic mice expressing the human ACE2 receptor (hACE2-Tg). Finally, our immunization strategy induced strong effector memory response, potentiating protective immunity against re-exposure to SARS-CoV-2 spike protein. Our results show the potential of a new Sindbis virus-based vaccine platform to counteract waning immune response that can be used as a new candidate to combat SARS-CoV-2. Given the strong T-cell responses elicited, our vaccine is likely to be effective against variants that are proving challenging, as well as, serve as a platform to develop a broader spectrum pancoronavirus vaccine. Similarly, the vaccine approach is likely to be applicable to other pathogens.

## 1 Introduction

In the ongoing COVID19 pandemic, vaccines play a key role in the strategy to bring SARS-CoV-2 transmission under control. Safety and eliciting a broad-spectrum immune response are paramount for coronavirus vaccine development. Data from vaccine clinical trials and real-world evidence show that available coronavirus vaccines are able to cut the risk of severe COVID19 disease and transmission. However, even with first generation vaccines currently being globally administered to reduce transmission and severity of the disease, the emergence of circulating variants has raised major concerns that challenge sustained vaccine efficacy, particularly in the face of waning immunity following vaccination[5; 6; 7; 8; 9; 10; 11]. Recent data have indicated that escape (appearance and spread of viral variants that can infect and cause illness in vaccinated hosts) protection by vaccines designed against the Wuhan-1 strain is inevitable[8].

The global COVID19 pandemic is unlikely to end until there is an efficient pan-global roll-out of SARS-CoV-2 vaccines. Though multiple vaccines are currently available, vaccine rollout and distribution at the time of writing this paper is quite incomplete. The three largest countries in the western hemisphere– US, Brazil, and Mexico – have vaccinated 32.7%, 7%, and 6.6% of their populations, respectively, compared to only 2.2% in India [12]. Vaccine distribution to date has been highly non-uniform among these and other countries around the globe, encountering many challenges. Unequal vaccine roll-out and the new B.1.617 variant are highly concerning. Major challenges have been supplies shortages, logistical problems, complex storage conditions, priced affordably, and safety[13]. Consequently, the pandemic is currently sweeping through India at a pace faster than ever before. The countries’ second wave became the worst COVID19 surge in the world, despite previous high infection rates in megacities that should have resulted in some immunity. More cost-effective and facilitated delivery of broad-spectrum SARS-CoV-2 vaccines would help improve wide and rapid distribution, which would in turn minimize vaccine-escape.

Traditionally, vaccines have been designed to induce antibody responses and have been licensed on their capacity to induce high titers of circulating antibody to the pathogen[1]. With increased knowledge of host-virus interactions, it has become clear that the cellular arm of the immune response is also crucial to the efficacy of vaccines against pathogens and to provide appropriate help for antibody induction. Various strategies have emerged that specialize in developing candidate vaccines that solely induce either cellular or humoral responses[1]. However, as most viruses and pathogens reside at some point during their infectious cycle in the extracellular as well as intracellular space, vaccines need to promptly elicit a strong T-cell memory response against intracellular pathogens, so that, at the earliest stages of the infective process, preventing disease can be addressed in coordination with antibodies.

It has been reported that recovered COVID19 patients consistently generate a substantial CD4+ T (OX40+CD137+) cell response against SARS-CoV-2 spike[3]. SARS-CoV-2-specific CD4+ T-cells produced IL-2 and substantial amounts of IFNg, hallmarks of Th-1 type effector T-cell polarization. Th-1 type effector T-cells provide critical help for CD8 T-cell priming and conferring cytotoxic T-cell mediated immune protection. The costimulatory molecule OX40 is a member of TNF receptor superfamily (TNFRSF) that is upregulated on activated T-cells shortly after T-cell receptor recognition of specific antigen[15; 16]. It is mainly expressed on CD4+ T-cells, although activated CD8 T also express OX40, albeit at lower levels[17]. Once activated, OX40 receptor is the key molecule for clonal expansion, differentiation and survival of Th1-effector cells and cytokine production. [15; 18; 19; 20; 21; 22]. Although OX40 does not directly initiate T-cell memory formation, it contributes to homeostasis of memory T-cells and enhances effector memory T-cell function[23]. In addition to its role in direct T-cell mediated viral clearance (T-cell immunity), OX40 stimulation is found to cooperate with the inducible costimulating (ICOS) molecule on follicular T helper (Tfh) cells augmenting their amplification and development to coordinate humoral immune response[24]. Antigen-specific activated Tfh cells help B cells produce high affinity antibodies against pathogens and are indispensable for vaccine induced long-lasting humoral immunity by facilitating differentiation of memory B cells and long-lived plasma cells from Germinal Centers (GC)[25; 26; 27]. Therefore, designing a vaccine that could stimulate OX40 would provide a powerful platform for T-cell mediate immunity.

Alphaviruses have demonstrated strong attributes as a development-and-manufacturing platform for vaccines[5; 6; 7; 8; 9; 10; 11]’[12], Particularly, studies with SARS-CoV strains bearing epidemic and zoonotic spike variants are promising[11]. The strength of the use of alphavirus vaccine utilization is the generation of rapid, high level, and transient nature of transgene expression [13]. Importantly, we have shown in our earlier preclinical work[29; 30; 31] that alphavirus vaccine platforms have the advantage to directly deliver antigens and immune modulatory molecules to lymph nodes, where they are expressed transiently to elicit diversified CD4+ and CD8+ T-cell immunity effective at controlling tumors throughout the body. These vectors represent a highly effective self-amplifying mRNA vaccine that can be engineered to express multiple antigens and stimulatory molecules. Within three hours after infection the vector generates hundreds of thousands of mRNA copies within the infected cells and high levels of expression of the transgenes (e.g., the spike antigen and anti-OX40 antibody). At the same time, the transient nature and cytosolic location of RNA improves the safety profile of SV vector-based vaccines. The replication defective nature of our vectors ensures no further transmission of the virus beyond the infected cells[14]. Replication-deficient alphavirus-based vaccines are immunogenic, safe, well tolerated and can be cost-effectively stored and transported using conventional 2-8 °C storage as well as lyophilization.

Here we describe a new Sindbis Virus (SV) vaccine transiently expressing the SARS-CoV-2 spike protein (SV.Spike), which induces a strong adaptive immunity that fully protects transgenic mice that express the SARS-CoV receptor (human angiotensin-converting enzyme 2 [hACE2]), hACE2-Tg, against authentic SARS-CoV-2 virus infection. In addition, we demonstrate that combination of our vaccine with αOX40 agonistic antibody significantly enhances the induction of immunity by the SV.spike vector. Specifically, seroconversion and abundance of IgG neutralizing antibodies and T-cell immunity through early initiation of Th1-type T-cell polarization are markedly augmented to potentiate long-term immunity protective against SARS-CoV-2 infection in mice. Together these studies develop a safe and effective vaccine platform that provides humoral and cellular immunity to the SARS-CoV-2 spike. This platform has the potential to be applied to other emerging pathogens.

## 2 Results

### 2.1 Construction and characterization of Sindbis carrying the SARS-CoV-2-spike

We designed and generated a Sindbis alphavirus replicon carrying the SARS-CoV-2 spike mRNA. SV vectors are generated from two plasmids: a replicon and helper (Figure1 and Supplementary Figure 1). Genes of interest (GOI) can be substituted for the 5kb structural genes that were removed to generate the helper plasmid. The plasmid encoding the structural genes does not contain a packaging signal, preventing further virus assembly beyond the initial preparation of the vectors in BHK-21 cells. Plasmids are transcribed from the T7 promoter and the RNA transcripts are electroporated into BHK-21 cells to produce viral vectors.

**Figure 1.**
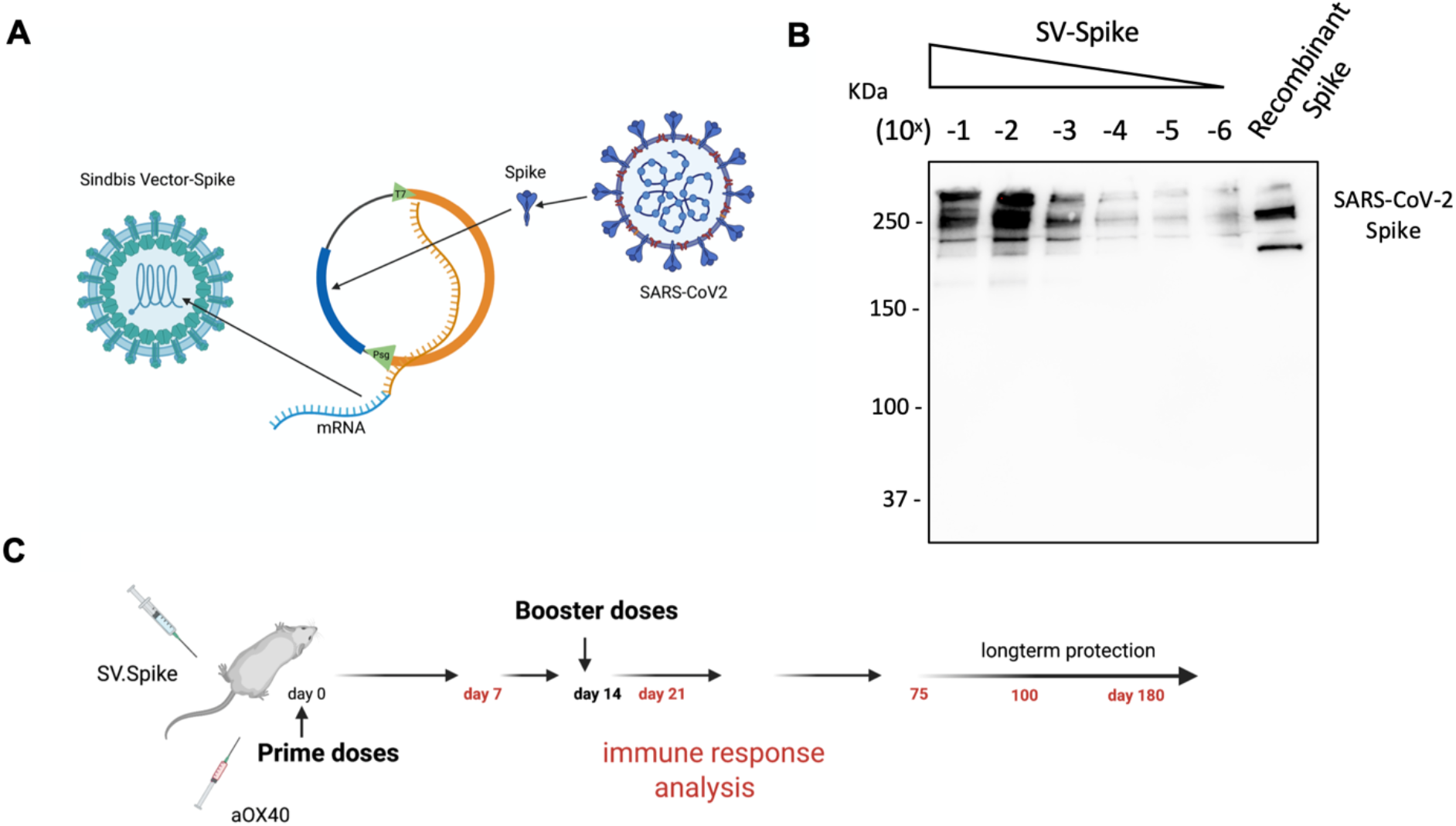
Characterization of Sindbis vector carrying the SARS-CoV-2 spike. **(A)** Schema of SARS-CoV-2 spike gene cloned into Sindbis vector system. **(B)** Western Blot of SARS-CoV-2 spike produced from the Sindbis vector. Lanes shown are titration of the vector, and recombinant spike control produced in HEK cells. **(C)** Schematic of vaccination. C57BL/6 mice were immunized with 1x 0.5 ml SV.Spike/and or αOX40 antibody (250μg/dose) on day 0. A boost injection of SV.Spike/and or αOX40 were once given on day 14. On day 7,14 and 21, 75 and 100, blood was taken to determine Sars-Cov-2 spike specific antibodies by ELISA. Spleens were excised and a single cell suspension was stained and analyzed by flow cytometry. T-cells were isolated and were used for ELISPOT assay and Seahorse. As control, naïve C57BL/6J mice were used.

The combination of SV vectors encoding a selected antigen with immunomodulatory antibodies makes them far more effective than they are alone[40; 41; 42]. In particular we have found that combining SV vectors expressing specific antigens with αOX40 generates very potent immune responses capable of eradicating tumors in multiple murine models and conferring long-term protection against tumor recurrences or rechallenges[40].

The overall design in the production of Sindbis SARS-CoV-2 spike (SV.Spike) is illustrated in Figure 1 and Supplementary Figure 1. We determined the expression of the full-length SARS-CoV-2 spike from infected cells by western blot in Figure 1B.

The immune responses induced by the Sindbis SARS-CoV-2 spike (SV.Spike) vaccine candidate were analyzed in C57BL/6J mice. Groups of mice (*n* = 5) were immunized by intraperitoneal (i.p.) route, by prime-boost vaccine strategy with SV.Spike and/or αOX40, with 14 days difference between the two doses (Figure 1C). Activation and priming of T-cells were analyzed by flow cytometry and ELISPOT at day 7, 21 post-immunization (p.i.), while cytotoxic assay and transcriptomic analysis was performed in T-cells isolated at day 7 p.i.. Metabolic activation of T and B cells was tested by Seahorse measurements (Agilent, CA) at day 7 and 21, respectively. Long-term memory T-cell analysis was carried out at day 100 p.i.. The overall antibody responses were measured at all the indicated time points (from day 7 to day 100 p.i.; Figure 1C).

### 2.2. Sindbis vaccine-elicited antibodies to SARS-CoV-2 spike

Serum IgM, IgG and IgA responses to SV.Spike, SV.Spike+αOX40, injections were measured on days 21, 75 and 100 days after vaccination by enzyme-linked immunosorbent assay (ELISA) against recombinant SARS CoV2 spike protein[3; 4]. Sera from all of mice tested showed reactivity to recombinant SARS-CoV-2 spike protein and, as might be expected, levels of antibodies varied based on the experimental group and time point. Consistent with previous reports[43; 44; 45], levels of IgM and IgG measured at day 21 and 75 post injection (p.i.) were significantly higher in the mice vaccinated with SV.Spike and combination of SV.Spike+αOX40 than in the mice who had received αOX40 alone or the naïve group (Figure 2A). Moreover, the SV.Spike+αOX40 group showed higher titers of IgG compared with only SV.Spike treatment, for which IgM was the predominant isotype and did not show seroconversion to IgG over the different time points. Specifically, both SARS CoV2-specific IgG and IgM antibodies demonstrated the highest expression on day 21 post immunization for the indicated groups (IgG-OD450 of 2.3 for SV.Spike+αOX40 serum, and IgM-OD450 of 1.9 for SV.Spike serum). At days 75 p.i., IgG were still significantly predominant in the sera of the mice immunized with the SV.Spike+αOX40 combination (IgG-OD450 = 1.3), whereas IgM reactivity did not significantly vary from day 21 to day 100 compared with the control groups (Figure 2B). Instead, IgM levels in the SV.Spike mice showed a more significant decrease and less lasting reactivity from days 21 to 75 days p.i. (IgM-OD450 of 1.2) compared to the control group, whereas the IgG trend demonstrated significant high reactivity only at day 21 p.i.. Conversely, IgA levels did not show any significant difference in any of the groups and time points tested (Figure 2A, B). These data support the evidence that immunization of mice with SV.Spike combined with αOX40 elicits a strong and specific immune response, which is predominantly represented by SARS-CoV-2 IgG- specific antibodies.

**Figure 2.**
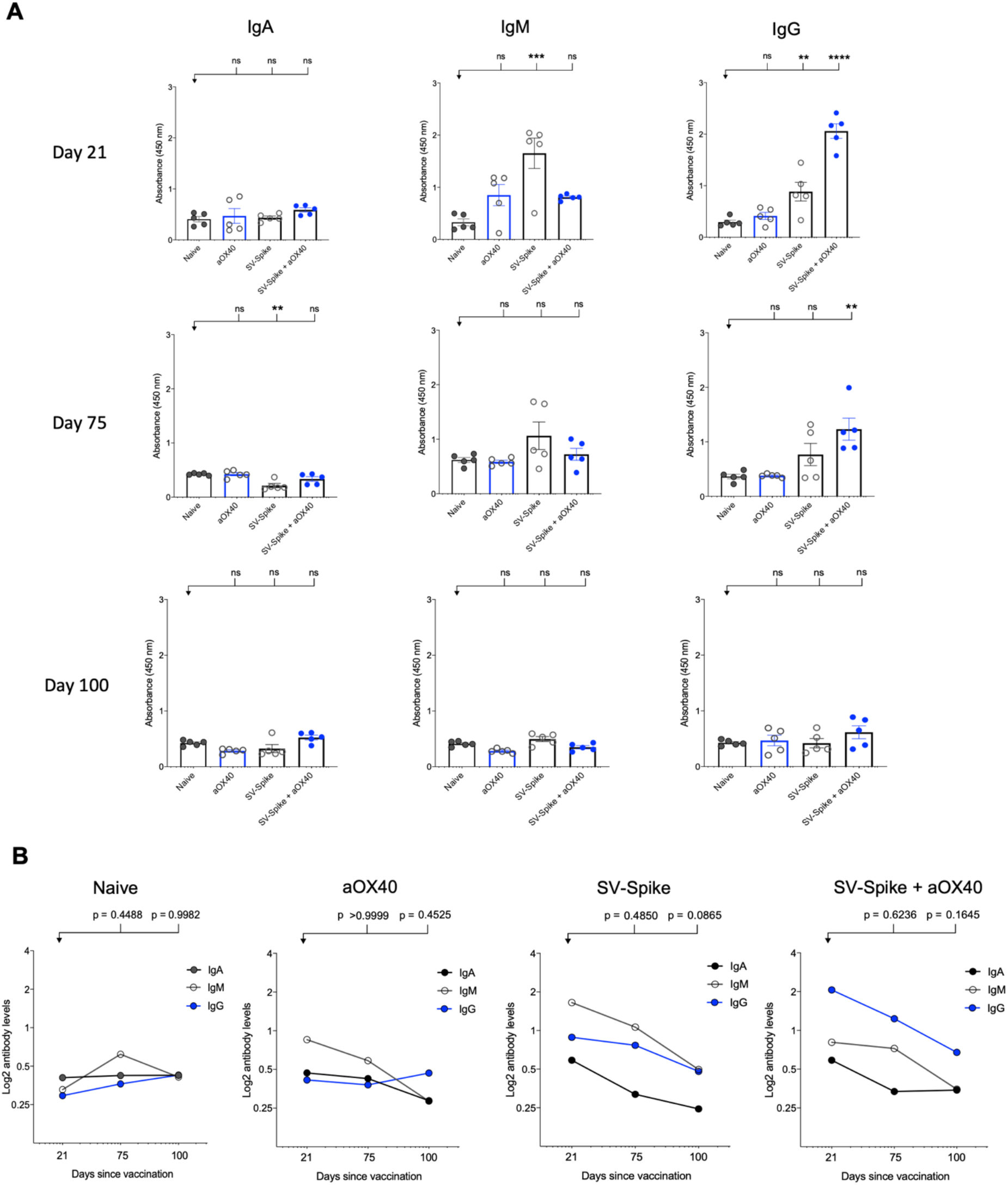
SARS-CoV-2 spike specific antibodies induced by Sindbis vector. Characterization of serum IgA, IgM, and IgG in C57BL/6J mice vaccinated with SV.Spike at day 21, 75 and 100 post-immunization. **(A)** The levels of Spike-specific IgA, IgM, and IgG isotypes in sera of immunized mice at different time windows. P values were calculated by one-way ANOVA with the Bonferroni correction in Graphpad Prism. n.s. > 0.05; **P < 0.01; ***P < 0.001; ****P<0.0001. **(B)** The kinetics of Spike-specific IgA, IgM, and IgG isotypes in sera of immunized mice at different time windows. Two-way ANOVA with the Bonferroni correction in GraphPad Prism used to calculate the indicated P values. The data presented are the mean of three technical replicates. The median values of **(A)** OD450 or **(B)** calculated log2 antibody levels were plotted for each isotype of three antibodies.

### 2.3 Anti-SARS-CoV-2 spike neutralizing antibodies induced in Sindbis vaccinated mice block the SARS-CoV-2 spike protein from binding to hACE2 receptor proteins

Immediately after SARS-CoV-2 was identified as the causative agent of the COVID-19 outbreak, it was shown that human ACE2 (hACE2) is the main functional receptor for viral entry[46]. We hypothesized that the virus–receptor binding can be mimicked *in vitro* via a protein–protein interaction using purified recombinant hACE2 and the Spike of the SARS-CoV-2 protein. This interaction can be blocked by virus naturalizing antibodies (NAbs) present in the test serum of vaccinated mice.

A competition ELISA assay was developed to detect whether SARS-CoV-2 spike-specific antisera from mice immunized with αOX40, SV.Spike and SV.Spike+αOX40 could block the interaction between SARS-CoV-2 spike and hACE2. Our assay demonstrated that the specific Spike–hACE2 binding can be neutralized by SV.Spike or SV.Spike+αOX40 sera in a dose-dependent manner, but not by sera from αOX40 alone or naïve groups (Supplementary Figure 2A, B). Similar results are obtained by the intramuscular route (Supplementary Figure 2C). As shown in Figure 3A, antibodies in the antisera from mice immunized with SV.Spike and combination of SV.Spike and αOX40 at day 21 post-immunization significantly inhibited the binding of SARS-CoV-2 spike to hACE2 compared to the sera from naïve mice, indicating that SV.Spike-induced antibodies could strongly neutralize SARS-CoV-2 infection by blocking the binding of Spike protein on the surface of SARS-CoV-2 to hACE2.

**Figure 3.**
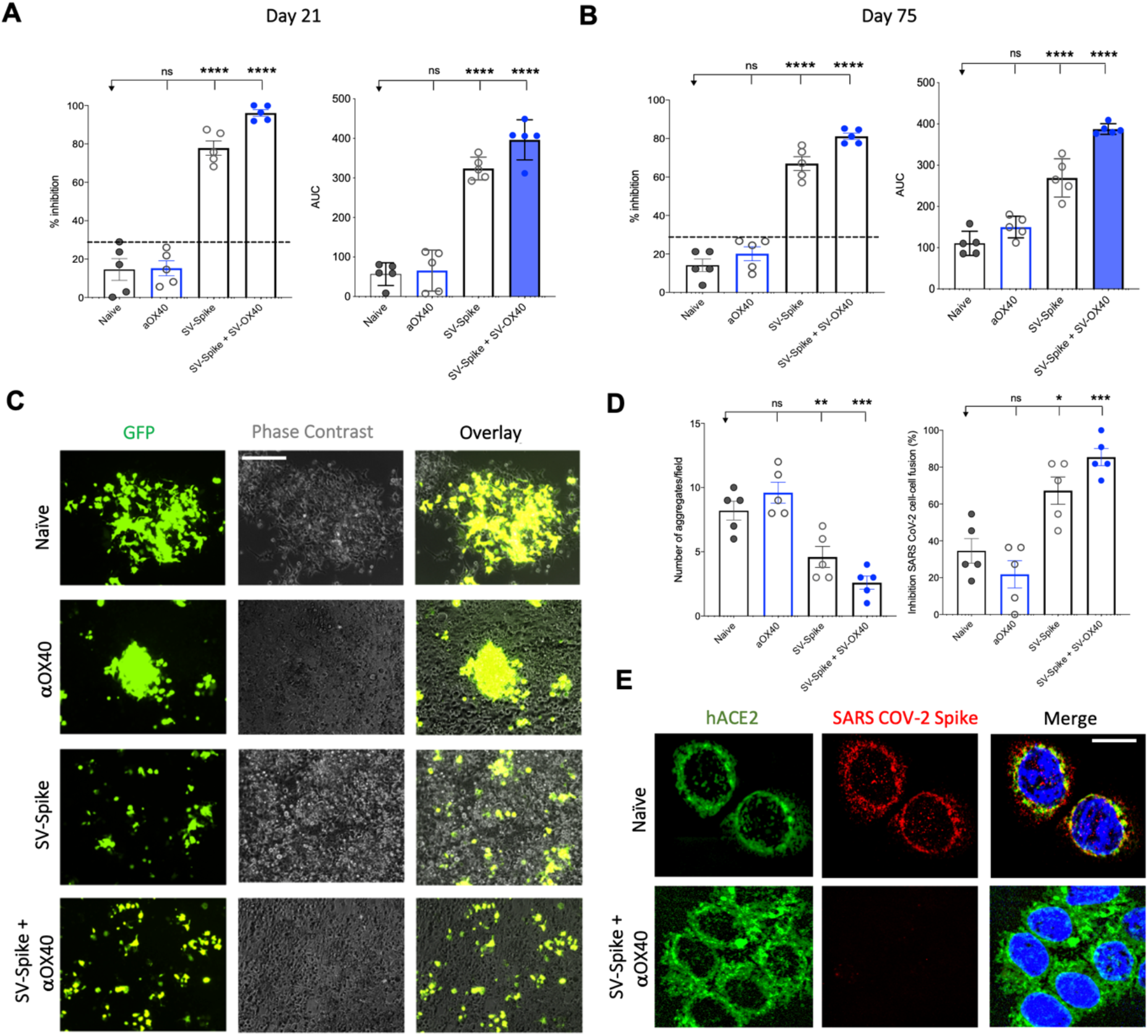
Blockade of SARS-CoV-2 spike-hACE2 binding and spike protein-mediated cell–cell fusion by anti-SARS-CoV-2 spike neutralizing antibodies. **(A, B)** In the assay, anti-SARS-CoV-2 neutralizing antibodies from immunized C57BL/6J mice, block recombinant Spike protein from binding to the hACE2 protein pre-coated on an ELISA plate. Percentage of inhibition distributed along y-axis of SARS-CoV-2 spike–hACE2 interaction for the indicated reciprocal plasma dilutions by mouse sera collected at **(A)** 21 and **(B)** 75 days post vaccination with Sindbis expressing SARS-CoV-2 spike (SV.Spike), SV.Spike in combination with αOX40 and αOX40 alone compared to the naive group. Area under the curve (AUC) values of serum antibodies were calculated from reciprocal dilution curves in antibody detection assay. The data presented are the mean of 5 biological replicates with two technical replicates. Statistics were performed using a One-way ANOVA with the Bonferroni correction in Graphpad Prism. n.s. > 0.05; *P < 0.05; **P < 0.01; ***P<0.001; ****P<0.0001. **(C)** Images of SARS-CoV-2 spike-mediated cell–cell fusion inhibition on 293T/ACE2 cells by sera from C57BL/6J vaccinated mice. SARS-CoV-2 spike-transfected 293T were incubated with mice serum at 1:100 dilution and applied onto 293T/ACE2 cells for 24 hours. Scale bar: 100 μm. **(D)** Quantification of the number aggregates (left panel) and inhibition of cell–cell fusions (right panel) induced by SARS-CoV-2 spike following pre-incubation with naïve, SV.Spike, SV.Spike+αOX40 and αOX40 alone are shown. N = 5 biological replicates with 2 independent technical replicates. One-way ANOVA with Bonferroni correction *P < 0.05, **P < 0.01, and ***P < 0.001. **(E)** Representative confocal images of 293T/ACE2 cells treated with serum from Naïve and SV.Spike+αOX40-immunized mice pre-incubated with SARS-CoV-2 spike recombinant protein and stained for hACE2 (green), SARS-CoV-2 spike (red), and DAPI (blue). Scale bar: 20 μm.

To investigate whether the neutralizing antibody response in immunized mice could maintain a high level for a longer period of time, we tested the neutralization activity of mice sera at 75 days post-immunization. The results showed that, although the overall antibody neutralizing capacity decreased compared to day 21, antibodies from SV.Spike and SV.Spike+αOX40 groups still significantly competed for the binding of the SARS-CoV-2 spike and hACE2 (Figure 3B), indicating that our SV.Spike vaccine is able to induce relative long-term neutralizing antibody responses.

Next, we investigated if the serum from mice immunized with SV.Spike could inhibit the cell membrane fusion process for viral entry[47; 48; 49]occurring upon the binding of SARS-CoV-2 spike Receptor Binding Domain (RBD) fragment to the ACE2 receptor on target cells. To establish an assay for measuring SARS-CoV-2-spike-mediated cell–cell fusion, we employed 293T cells (a highly transfectable derivative of human embryonic kidney 293 cells, that contain the SV40 T-antigen) expressing both SARS-CoV-2 spike and enhanced green fluorescent protein (EGFP) as effector cells and 293T cells stably expressing the human ACE2 receptor (293T/ACE2) as target cells. Notably, when the effector cells and the target cells were co-cultured at 37°C for 6 h and 24 h, the two types of cells started to fuse at 6 h, exhibiting a much larger size and multiple nuclei compared to the unfused cells. These changes were more significant at 24 h, resulting in hundreds of cells fused as one large syncytium with multiple nuclei that could be easily seen under both light and fluorescence microscopy (Supplementary Figure 3). The cell fusions were observed in the cells transfected with SARS-CoV-2 spike but not SARS-CoV Spike, whereas those cells transfected with EGFP only did not elicit such an effect, confirming that CoV-2 Spike-hACE2 engagement is essential for viral fusion and entry.

To determine whether the serum of mice immunized with SV.Spike can block Spike protein-mediated cell–cell fusion, we incubated the effector cells with serum from Naïve, SV.Spike and/or αOX40 mice (diluted 1:100) at 37 °C for 1 h and then we co-cultured them with the 293T/ACE2 target cells. We found that not only were fewer fusing cells observed, but also the size of fused cells were visually smaller in the groups of SARS-CoV-2-spike/293T effector cells pre-incubated SV.Spike with or without αOX40 sera compared to controls (Figure 3C). Quantification of fused cells per field in at least four randomly selected fields revealed a remarkably lower number of cell–cell fusions in both SV.Spike and SV.Spike+αOX40 groups compared to all the other groups. Moreover, SARS-CoV-2 spike-mediated cell–cell fusions were significantly inhibited by serum derived from SV.Spike+αOX40 vaccinated mice, indicating that addition of αOX40 to the vaccination protocol elicits antibodies with enhanced interference of syncytium formation mediated by SARS-CoV-2 infection (Figure 3C, D).

The interference of immunized sera NABs with SARS-CoV-2-hACE2 binding was also determined by immunofluorescence experiments performed by culturing 293T/ACE2 cells with recombinant SARS-CoV-2 spike previously incubated with serum from naïve and SV.Spike and αOX40 immunized mice. The binding between Spike and hACE2 expressed on the cell surface was subsequently visualized via confocal fluorescence microscopy (Figure 3E). As expected, Spike incubated with SV.Spike+αOX40 serum was incapable of binding to hACE2, while the control group showed evident co-localization with hACE2 on the cell surface.

Taken together, these data demonstrate that SV.Spike alone and to a greater extent SV.Spike+αOX40 sera can neutralize SARS-CoV-2 spike-hACE2 interaction and in turn counteract virus entry mediated by cell-membrane fusion.

### 2.4. SV.Spike vaccine prevents infection of SARS-CoV-2 in transgenic hACE2-Tg mice

The neutralizing activity of serum from vaccinated mice was determined using Luciferase-encoding SARS-CoV-2 spike pseudotyped lentivirus[50; 51] [52] (Supplementary Figure 5A, C), by testing the impact of the serum on the lentivirus transduction. Serial dilutions (1:300, 1:600, 1:900: 1:1800, 1:3200 and 1:6400) of mice sera harvested at day 21 and 75 p.i. were incubated with equal amounts of lentivirus for 1 hour at 37 °C, then plated on 293T/ACE2 cells. We then measured the amount of blocked pseudotyped viral particles in infected cells by determining the amount of luminescence reduction, which reflects the level of neutralizing antibody or molecular inhibitors in the sample. The results showed that the antisera could inhibit SARS-CoV-2 pseudotype infection in a dose-dependent manner (Supplementary Figure 5), consistent with the result from the antibody neutralization assay (Supplementary Figure 3). Our results demonstrate that sera from SV.Spike with or without αOX40 immunized mice groups resulted in significantly high levels of neutralizing antibodies both at day 21 and 75, since they overcame the pseudotyped lentivirus infectivity inhibition threshold of 30% (Figure 4A, B). Moreover, serum from these mice receiving combination of SV.Spike and αOX40 gave the highest levels of neutralization at day 21 after vaccination (95.3% of inhibition), with a slight decrease at day 75 (79% of inhibition). Naïve and αOX40 groups did not develop a neutralizing antibody response (% inhibition < 30%) at the timepoints tested, consistent with their lack of SARS-CoV-2 spike binding antibodies.

**Figure 4.**
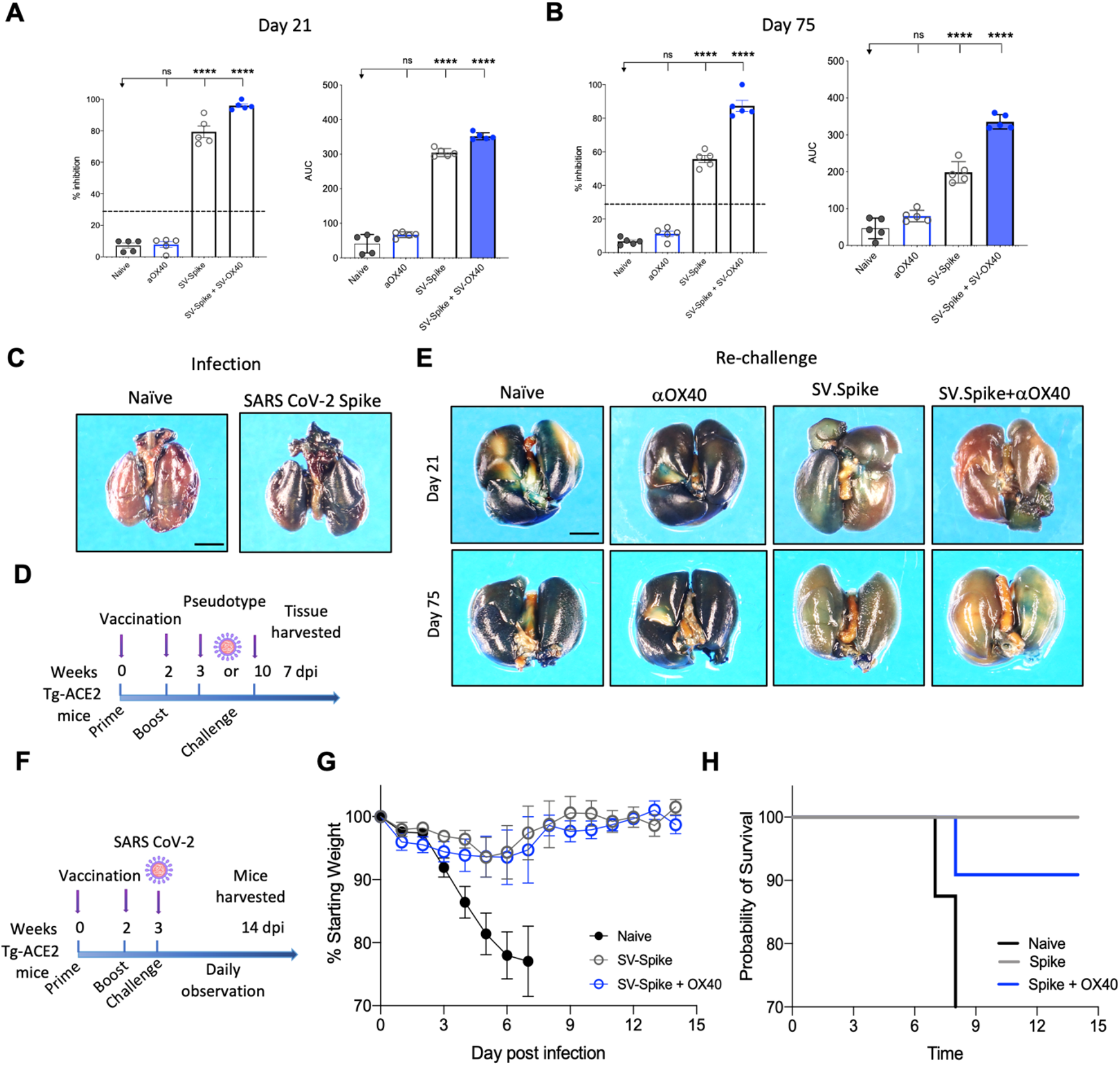
Sindbis-Spike vaccine prevents infection of SARS-CoV-2 in hACE2 transgenic (hACE2-Tg) mice. *Luciferase*-encoding SARS-CoV-2 spike pseudotyped lentivirus was incubated with mouse sera collected at **(A)** 21 and **(B)** 75 days post vaccination with SV.Spike, SV.Spike in combination with αOX40 and αOX40 antibody alone compared and unvaccinated naïve groups. Area under the curve (AUC) values of serum antibodies were calculated from reciprocal dilution curves in antibody detection assay. The data presented are the mean of 5 biological replicates with two technical replicates. Statistics were performed using a One-way ANOVA with the Bonferroni correction in GraphPad Prism. n.s. > 0.05; ****P<0.0001. **(C)** Expression of pseudotyped SARS-CoV-2-spike-lacZ lentivirus in whole mouse lung following intranasal delivery. One week following vector nasal administration to the right nostril of four weeks old hACE2 transgenic mice (B6(Cg)-Tg(K18-ACE2)2Prlmn/J), expression of lacZ was analyzed in mice airways. X-Gal stained whole lungs from (left) hACE2 non carrier control mouse and (right) hACE2 transgenic mouse, both dosed with SARS-CoV-2-spike-lacZ pseudotyped lentivirus. **(D)** Schematic of the re-challenge experiment with SARS-CoV-2-spike-lacZ lentivirus. **(E)** On day 21 (upper panels) and 75 (lower panels) after the initial infection hACE2-Tg were rechallenged with 3.6 x 10^5^ PFU of SARS-CoV-2-spike-lacZ pseudotyped lentivirus and then analyzed for X-Gal staining at day 7 post rechallenge. Three non-vaccinated naïve animals were included as a positive control in the rechallenge experiment. **(F-H)** hACE2-Tg mice were vaccinated with SV.Spike and/or αOX40 and challenged with 10^4^ particles of live SARS-CoV-2 coronavirus at day 21 post immunization. Weight loss and mortality was observed daily for 14 days after live virus infection and compared to the naïve unvaccinated group. **(G)** Change of body weight during systemic infection with SARS-CoV-2 coronavirus. Percent weight loss (y-axis) is plotted versus time (x-axis). Data points represent mean weight change +/- SEM. **(H)** Survival curves of SV.Spike with or without αOX40 treated and naïve unvaccinated mice. n = 5 mice per group.

Recently, hACE2 transgenic (B6(Cg)-Tg(K18-ACE2)2Prlmn/J or hACE2-Tg) mice were used for the development of an animal model of SARS-CoV-2 infection[53]. In order to test pseudotyped lentivirus infectivity rate *in vivo,* we produced a *nLacZ*-encoding lentivirus expressing SARS-CoV-2 spike protein (Supplementary Figure 4B, D) and we evaluated the vector expression following delivery to hACE-Tg mice airways, by administrating a single dose of *nLacZ*-pseudotype to 4-week-old hACE2-Tg mice by intranasal inhalation. After 7 days, the airways were harvested and intact glutaraldehyde-fixed tissues were processed for staining with X-Gal for detection of β-galactosidase activity expressed from the nuclear-localized lacZ reporter gene (*nlacZ;* Figure 4C). Positive X-Gal staining observed in airways upon lentivirus intranasal administration indicated the successful SARS-CoV-2-spike lentiviral vector expression and pseudotype delivery in mice airways.

In order to investigate the protective effects of SV.Spike vaccination *in vivo,* we subsequently immunized hACE2-Tg mice with the same strategy as used for the C57BL/6J mice (Figure 1D). The hACE2-Tg mice were vaccinated at 0 and 2 weeks and then challenged with pseudotyped SARS-CoV-2 intranasally at day 21 and 75 post-immunization (Figure 4D). The lungs were collected at 7 days post-challenge and pseudotype delivery was tested by X-Gal staining. As shown in Figure 4E, the *nLacZ*-SARS-CoV-2-spike lentivirus could not be detected in the lungs from SV.Spike+αOX40 immunized mice, while substantially reduced infectious virus burden was still detected in the lungs from SV.Spike treated mice compared with the naïve group at the indicated time points. As expected, lungs from animals treated with αOX40 showed high amount of pseudotype particles, as indicated from the very high signal of X-Gal staining (Figure 4E). Finally, protective immunity was also assessed in young adult vaccinated Tg-ACE2 mice challenged with live SARS-CoV-2 coronavirus. Three weeks after prime and boost vaccination doses, all mice were challenged with 10^4^ particles of SARS-CoV-2 via the intranasal (i.n.) route (Figure 4F). We recorded the daily the body weight of each mouse after infection for a total of 14 days and found that the body weights of both SV.Spike and SV.Spike+αOX40 mice showed a slow decrease at 3-5 days post infection (dpi), with a progressive stabilization and increase of their weight at day 8-9 post infection. The naïve group showed a faster decrease during 3–5 dpi (Figure 4G), which led to early mortality around day 8 dpi (Figure 4H). Vaccinated mice did not evidence any signs of disease at the time the experiment was terminated but were culled on day 14 as required by the protocol, which was performed in an ABSL3 facility. Together, these data suggest that combination of SV.Spike and αOX40 vaccine in mice conferred remarkably long-term protection against SARS-CoV-2 infection by eliciting a durable humoral response in mice.

### 2.5 SV.Spike in combination with αOX40 metabolically reprograms and activates T-cells shortly after prime vaccine doses

Analysis of SARS-COV-2 specific adaptive immune responses during acute COVID-19 identified coordination between SARS-COV-2-specific CD4+ T-cells and CD8+ T-cells in limiting disease severity[54]. We analyzed vaccine elicited T-cell responses in the spleen 7 days after mice received prime doses of SV.Spike and/or αOX40 and compared the initial T-cell response to naïve mice (Figure 5). Spleens of mice were excised and a single cell suspension was stained and analyzed by flow cytometry.

**Figure 5.**
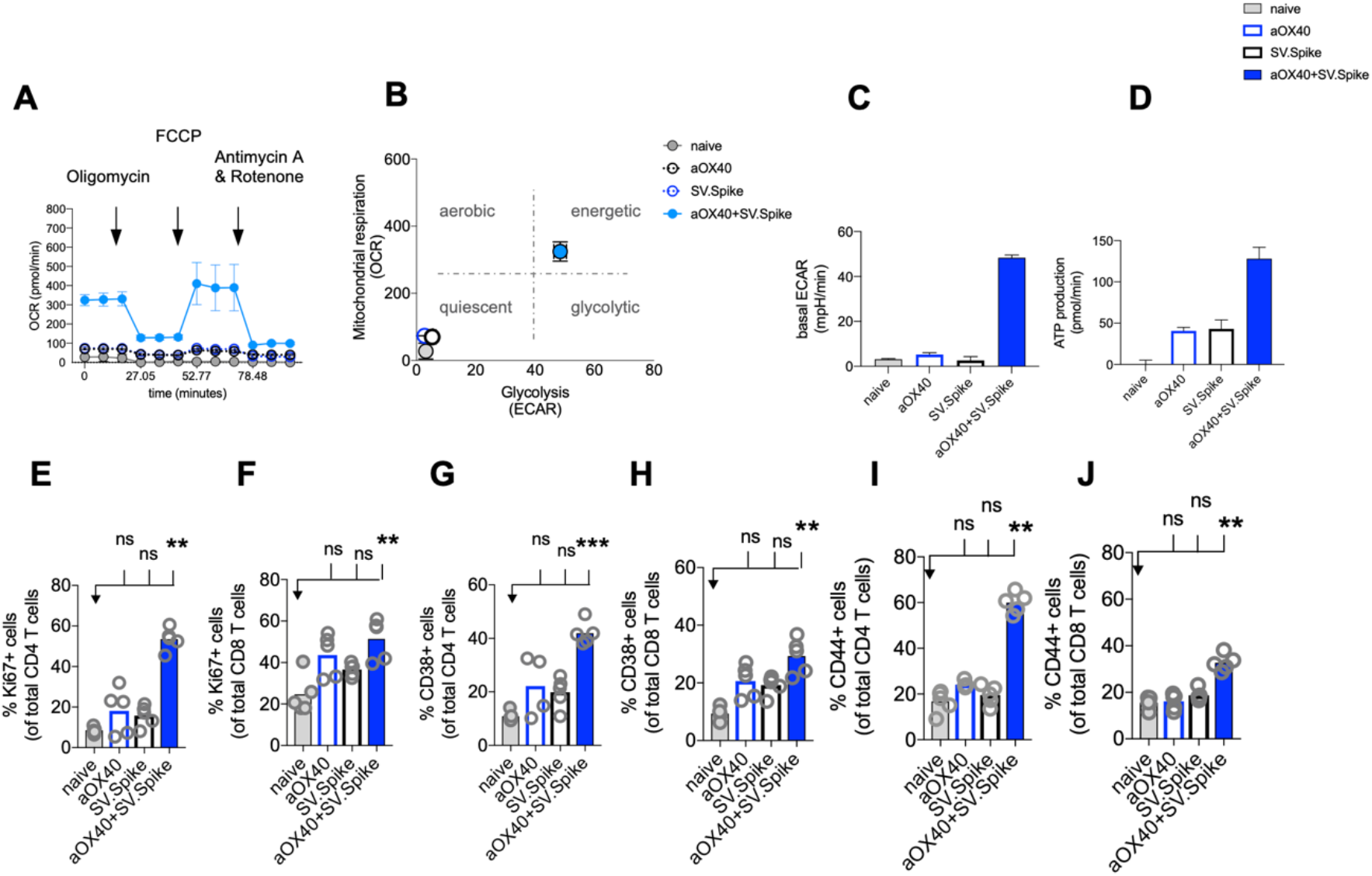
SV.Spike in combination with αOX40 activates and metabolically reprograms T-cells. C57BL/6J mice were vaccinated with first doses of SV.Spike and/or αOX40. Naive mice were used as control. T-cells were isolated from spleens on day 7 or otherwise indicated. **(A)** Mitochondrial respiration was assessed by measuring the median values of oxygen consumption rates (OCR) in T-cells of indicated groups using an extracellular flux analyzer. Oligomycin, FCCP, Antimycin A and Rotenone were injected as indicated to identify energetic mitochondrial phenotypes. **(B)** Energy Map (OCR versus ECAR) of T-cells from naïve or mice treated with SV.Spike, or αOX40 or combination of SV.Spike+αOX40 on day 7. **(C)** Baseline extracellular acidification rates (ECAR) in T-cells of indicated groups. **(D)** ATP Production in T-cells of indicated groups. (E-J) Splenocytes were analyzed in flow cytometry. **(E, F)** Expansion of CD4+ **(E)** and CD8+ T **(F)** cells is indicated by expression of Ki67-positive cells. **(G, H)** Activation of CD4+ T-cells **(G)** and CD8+ T-cells **(H)** indicated by CD38+ expression. **(I, J)** Expression of CD44+ positive cells. CD4 **(I)** and CD8 **(J)** cells. Error bars indicate SEM. Results are representatives of two independent experiments. Each symbol represents an individual mouse in E, F, G, H, I, J. Bars represent means. Statistical significance was determined with the Kruskal-Wallis test followed by the Dunns’ test. n.s. > 0.05, **p<0.005, ***p≤ 0.001.

For a successful vaccine-elicited immune response, differentiation of virus-specific T-cells from the naïve to the effector state requires a change in the metabolic pathways utilized for energy production[55]. Therefore, metabolic profiles of vaccine-induced T-cells are of interest and correlate to vaccine-mediated immunity[56].

We performed metabolic analysis of isolated T-cells from spleens in an Extracellular Flux Analyzer XFe24 (Seahorse Bioscience) to investigate metabolic changes of T-cells. We found, that combining our SV.Spike vaccine with agonistic αOX40 antibody metabolically rewires T-cells *in vivo* shortly after initial vaccine doses (Figure 5A-D). T-cells freshly isolated from mice on day 7 after first doses with SV.Spike+αOX40 combination displayed a metabolic shift to a highly bioenergetic state compared to single agent treatment or naïve mice that show a quiescent metabolism (Figure 5A-B). Naïve T-cells are quiescent and characterized by a metabolic program that favors energy production over biosynthesis. Upon T-cell receptor (TCR)-mediated stimulation, T-cells become activated and metabolically reprogrammed. The bioenergetic state of metabolically reprogrammed T-cells is characterized by a strong increase of oxygen consumption rate (OCR), which is a parameter for mitochondrial respiration (Figure 5A), and a strong increase of baseline extracellular acidification rate (ECAR) (Figure 5C), which is measured as a parameter for glycolysis. It has been shown that TCR signaling is directly tied to glycolysis[57]. We found that T-cells isolated from mice vaccinated with SV.Spike+αOX40 displayed a 3-fold increase of OCR and a 10-fold increase of ECAR compared to naïve and single agent vaccinated mice. T-cells switched to the energetic state ramped up their ATP production (Figure 5D). A metabolic rapid adaptation is further required for effector T-cells cytokine production and signaling. Rapid switch to type-1 cytokine production, such as IFNg and granzyme B (GrB) in antiviral CD8+ T-cells is more reliant on oxidative phosphorylation[58]. Indeed, immunophenotyping of CD4+ and CD8+ T-cells by flow cytometry revealed rapid clonal expansion of CD4+ T and CD8+ T subsets within one week after prime vaccine doses indicated by Ki67 expression on gated CD4+ and CD8+ T-cells. CD4+ T-cells showed the highest expansion increase by 10-fold in the combination vaccinated group compared to naïve and SV.Spike and αOX40 single agent immunized mice (Figure 5E-F). Both T-cell subsets were highly activated, indicated by CD38 and CD44 expression (Figure 5G-J) underlining successful vaccine elicited effector T-cell engagement by our vaccine shortly after initial vaccine doses. Similar results were obtained by the intramuscular route (Supplementary Figure 6).

### 2.6 SV.Spike+αOX40 vaccinated mice are characterized by a unique T-cell transcriptome signature profile after prime vaccine doses

To reveal the molecular profile of SV.Spike vaccine induced T-cell responses, we isolated T-cells 7 days after prime vaccine doses from spleens of mice from SV.Spike and/or αOX40 vaccinated groups and naïve group. We then performed mRNA deep sequencing (RNAseq) and network analysis (Figure 6). Principal-component analysis (PCA) showed a distinct segregation between combined SV.Spike and αOX40 vaccination and all other groups (Figure 6A). These data suggest, that SV.Spike and αOX40 induces a distinct T-cell response. Indeed, we next looked at gene expression profiles of naïve versus SV.Spike and/or αOX40 and we found that naïve versus SV.Spike+αOX40 markedly showed the highest amount of uniquely upregulated and downregulated total genes with 1,126 upregulated (left) and 328 downregulated transcripts (Figure 6B). Overall, in all groups more genes were significantly upregulated than downregulated (Figure 6B-C). These data suggest that SV.Spike+αOX40 changes the transcriptome signature of T-cells. We performed Gene Ontology (GO) functional enrichment analysis (also Gene Set Enrichment Analysis, GSEA) and network analysis from naïve mice versus SV.Spike+αOX40 (Figure 6D) and naïve versus SV.Spike only (Figure 6E) immunized mice to determine key pathways and intersections of these pathways. The majority of pathways were upregulated in T-cells isolated from mice immunized with SV.Spike+αOX40 with the exception of one cluster downregulated (ribosomal biogenesis). The upregulated pathways in the combination immunized mice were dominated by immune response, T-cell activation, chemokine/cytokine signaling, immune cell migration, DNA replication, chromosomal organization, cell cycle regulation, and chromatin modification that formed the central nodes of this network (Figure 6D). SV.Spike single agent immunized mice showed a smaller network of seven upregulated pathways including a main cluster of immune response closely connected to a cluster for to B cell engagement, a small cluster of cytokine production, chemotaxis, cell cycle, DNA replication, regulation of ROS (Figure 6E).

**Figure 6.**
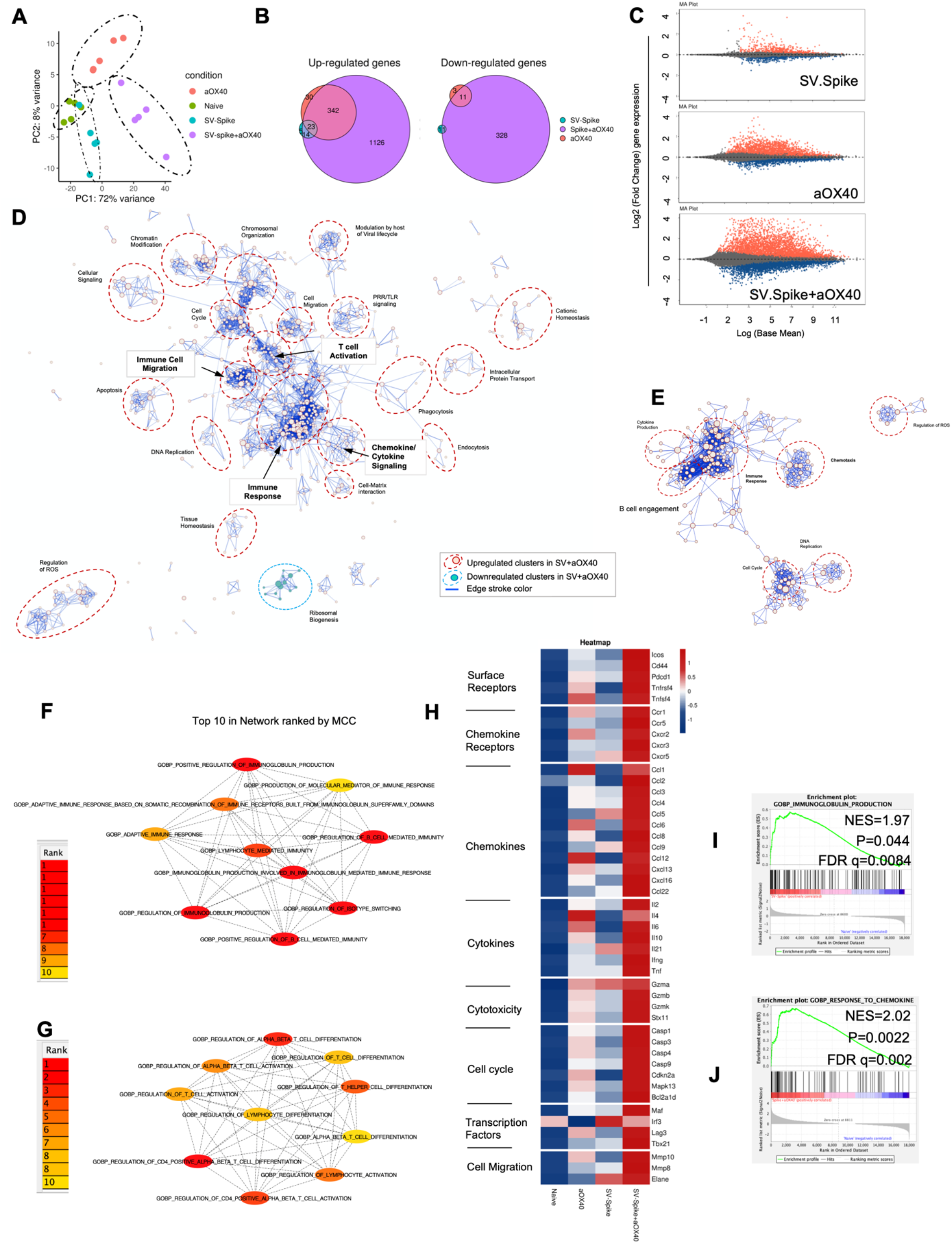
Sindbis expressing SARS-CoV-2 spike+αOX40 C57BL/6J vaccinated mice are characterized by a unique transcriptional signature of T-cells. Combination therapy markedly changes the transcriptome signature of T-cells favoring T-cell differentiation towards effector T-cells with a Th1 type phenotype 7 days after prime vaccination. **(A)** Principal component analysis (PCA) of RNA seq data from naïve, SV.Spike and/or αOX40 groups. **(B)** Venn diagrams summarizing the overlap between differentially expressed genes (DEGs) from SV.Spike (blue), αOX40 (pink) and SV.Spike+αOX40 (purple). Up-regulated DEGs (left) and down-regulated (right). **(C)** MA plots of differentially expressed genes in T-cells of naive versus SV.Spike (top graph), αOX40 (middle graph) and combination (bottom graph). Significantly (p<0.05) upregulated and downregulated DEGs are depicted in red or blue, respectively. **(D)** Pathway and network analysis based on GSEA in T-cells isolated from mice treated with combination therapy. Downregulated (blue circle) and upregulated (red circles) pathways are shown, respectively. **(E)** Pathway and network analysis based on GSEA in T-cells isolated from mice treated with single dose of SV.Spike. Top 10 hub biological process gene ontology (GO) terms ranked by the Cytoscape plugin cytoHubba (red, highest ranks; yellow, lowest ranks) in the SV.Spike only **(F)** versus combination immunized group **(G)**. Heatmap analysis of selected genes based on normalized read counts linked to T-cell differentiation in the SV.Spike and/or αOX40 immunized mice compared to naïve **(H)**. Highlighted selected gene set enrichment analysis (GSEA) pathways based on DEG in naive versus SV.Spike **(I)** and combination treated group **(J)**.

We next identified the top 10 hub GO terms by employing the Maximal Clique Centrality (MCC) for SV.Spike (Figure 6F) and SV.Spike+αOX40 (Figure 6G) immunized mice. We found that top 10 hub GO terms in SV.Spike only immunized mice were a selected network cluster of B cell stimulation and Immunoglobulin regulating pathways compared to the combination that represents a cluster of lymphocyte activation and differentiation regulating pathways. Additionally, we performed Protein Association Network Analysis using STRING to identify differentially expressed genes (DEGs)-encoded protein-protein interactions (PPIs). Significantly upregulated DEGs (log2FC>2, p<0.05) in T-cells of SV.Spike and/or αOX40 vaccinated mice compared to naïve were analyzed to assess overrepresentation of Gene Ontology (GO) categories in Biological Processes in all groups (Supplementary Figure 7). GO Biological Processes (Strength ≥1; p<0.05) identified by STRING for each group were assigned to one of 7 clusters (apoptosis, light green; cell cycle, red; cellular signaling, dark blue; chemokines/chemotaxis, yellow; cytokines, pink; immune response, light blue; mitochondrial ATP production, dark green). Each GO Biological Process term is defined by one gene set. The amount of contributing DEGs from mice immunized with SV.Spike and/or αOX40 in each gene set is shown as percentage. We identified fourteen biological processes for αOX40, thirteen for SV.Spike and forty-five for the combination vaccine strategy. We found cell-cycle related processes solely in the SV.Spike+αOX40 combination. The highest amount of chemokines/chemotaxis related processes was observed in the combination (eleven) compared to αOX40 (four) and SV.Spike (four) alone. Six cytokines related pathways were upregulated in the combination versus SV.Spike (one) and αOX40 (two) and fourteen immune response related terms were upregulated in the combination versus SV.Spike (four) and αOX40 (three). Overall, the percentage of DEGs that contribute to each biological process was highest in the combination vaccinated group compared to SV.Spike and αOX40 alone. Top 20 ranking of selectively enriched GO terms in the GSEA (FDA<0.05) revealed (GO) immunoglobulin production in the SV.Spike group (Figure 6H) and (GO) response to chemokine in the combination immunized mice group (Figure 6I, J). We analyzed expression of single signature gene transcripts for each immunized mouse group. We found the highest upregulation of DEGs (p<0.05) indicating T-cell dependent B cell stimulation for building up humoral immunity against SARS-CoV-2 (*ICos, Cxcr5, Il21, Cxcl13*), differentiation of Th-1 type effector T-cells associated with vaccine effectiveness (*Tnfrsf4, Cd44, ICos, Cxcr3, Ccr5, Il2, Ifng, Tbx21, Ccl3, Ccl4, Ccl9*) and antiviral cytotoxic T-cell stimulation for T-cell immunity (*Gzma, Gzmb, Gzmk*) in the SV.Spike+αOX40 immunized mice compared to single agent treated groups (Figure 6H).

In conclusion, these findings indicate that synergistic SV.Spike+αOX40 vaccine combination successfully changes the transcriptome profile of T-cells that is indispensable for building up humoral and T-cell immunity.

### 2.7 CD4+ T-cell help promotes effector differentiation of cytotoxic T-cells

SARS-CoV-2-specific T-cells are associated with protective immune responses[54]. Th1- type differentiated effector CD4+ T helper cells promote the development of CD8+ T-cells into anti-viral cytotoxic T lymphocytes (CTLs) and functional memory T-cells that can be quickly mobilized to directly kill SARS-CoV-2 early on upon re-infection preventing disease in coordination with SARS-CoV-2 specific humoral immune responses. CD4+ T helper cells are critical for success of vaccines and generally work by providing cytokines. We performed flow cytometry analysis to investigate CD4+ T helper differentiation, formation and antiviral cytotoxic effector T-cell differentiation in T-cells from SV.Spike and/or αOX40 immunized animals (Figure 7). Chemokine receptors help with the recruitment of type 1 effector and cytotoxic T-cells to tissues and lymphoid organs, site- specific activation of memory T-cells and T-cell clustering around activated antigen presenting cells (APCs). For example, virus-specific cytotoxic T lymphocytes (CTLs) are quickly recruited to influenza-infected lungs by a Th1 response, specifically due to the production of IFNg[59].Vaccines mimicking an infection can help to build up tissue specific immunity. Two of these Th1-type effector T-cell chemokine receptors are CXCR3 and CX3CR1. We found a significant increase of CXCR3 and CX3CR1 positive expressing CD4+ T-cells (Figure 7A, B) from spleens 7 days after administration of prime vaccine doses in the SV.Spike+αOX40 immunized mice group indicating effective recruitment and mobility of generated Th1-type effector T-cells. Immunophenotyping by flow cytometry revealed a 2-fold increase of the transcription factor Tbet and immune costimulatory molecule ICOS-double-positive Th1-type effector CD4+ T-cells compared with single agent vaccinated mice. Tbet+ ICOS+ are hallmarks of Th1-type T-cell polarization (Figure 7C, D).

**Figure 7.**
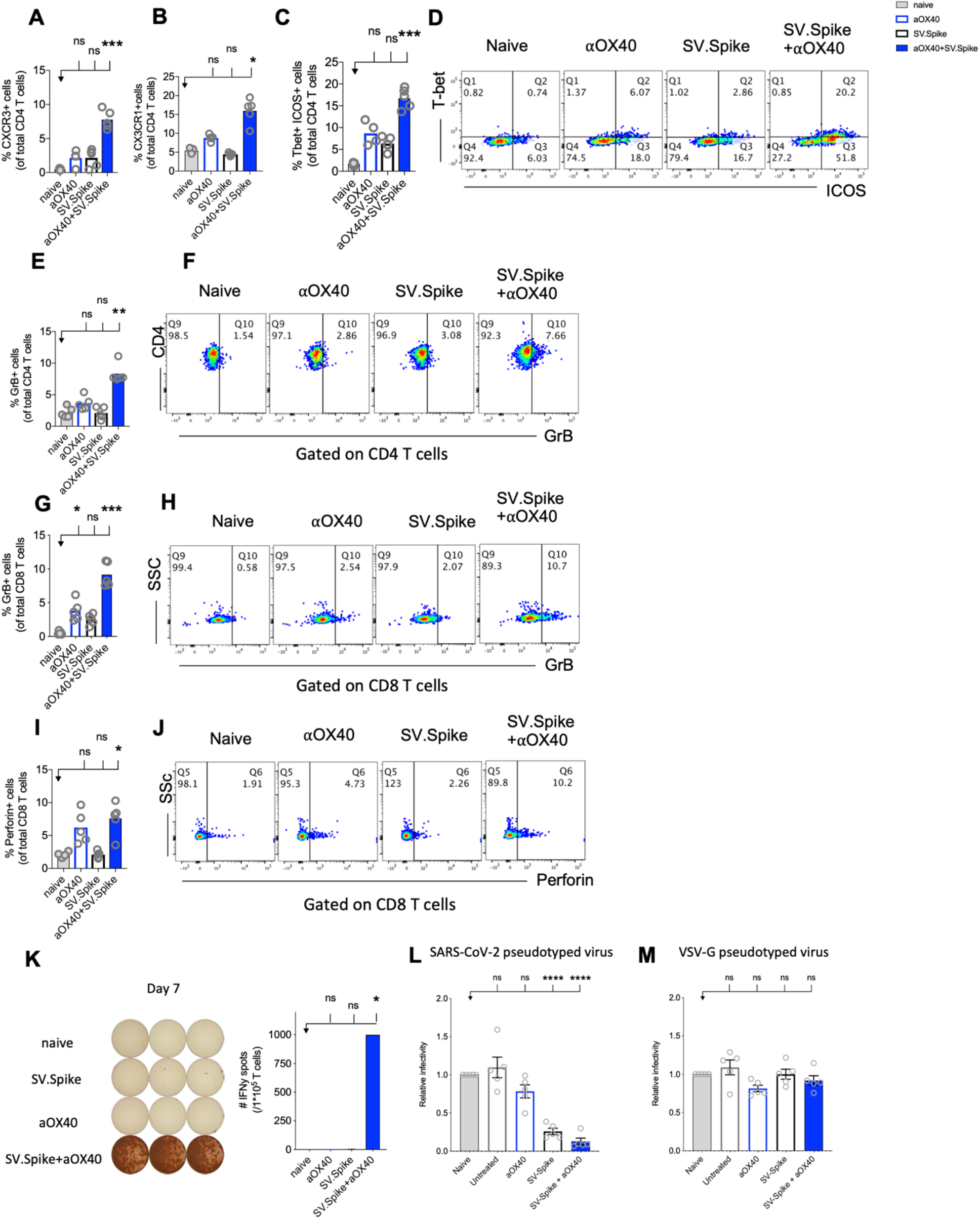
Reprogrammed T-cells in SV.Spike+αOX40 vaccinated mice display enhanced Th-1 T-cell phenotype mediated cytokine production and cytotoxic T-cell activity. Spleens of naïve and C57BL/6J vaccinated mice were excised on day 7 after prime vaccine doses for flow cytometry analysis **(A-J)**. T-cells were further isolated for **(K)** Interferon-g (IFNg) enzyme-linked immunospot analysis (ELISpot) and **(L, M)** cytotoxicity analysis. Percentage of **(A)** CXCR3 and **(B)** CX3CR1 expressing CD4+ T-cells indicating Th1-like T-cell effector phenotype. **(C)** Percentage of Tbet+ICOS+ positive Th1-type effector CD4+ T-cell polarization. (D) Representative blots. (E) Percentage of granzyme B (GrB) positive CD4+ T-cells from indicated groups using flow cytometry. **(F)** Representative blots. **(G)** Percentage of GrB positive CD8+ T-cells from indicated groups using flow cytometry. **(H)** Representative blots. **(I)** Percentage of Perforin positive CD8+ T-cells. **(J)** Representative blots. Bars represent means ± SEM (A-J) and each symbol represent an individual mouse (n=5 per group). Statistical significance was determined with the Kruskal-Wallis test followed by the he Dunns’ test. Results are representatives of at least two independent experiments. **(K)** Amount of IFNg spots per 10^5^ T-cells determined by ELISpot. **(L, M)** Cytotoxic activity of T-cells harvested on day 7 from control and treated mice (n = 5 mice per group). T-cells were isolated from splenocytes and were co-cultured with 293T/ACE2 cells for 2 days. Effector-to-target (E/T) cell ratio (T-cells/ACE2 cells) was 30:1. Cytotoxicity was determined for each group of mice by measuring the infectivity of luciferase-encoding pseudotyped particles with **(L)** Spike protein of SARS-CoV-2 or **(M)** VSV-G and is shown relative to naive T-cells. Bars or symbols represent means ± SEM, and statistical significance was determined with one-way ANOVA with the Bonferroni correction. n.s. > 0.05, *p<0.05, ***p≤ 0.001, ****p ≤ 0.0001.

The predominant pathway used by human and murine CD8+ T-cells to kill virus-infected cells is granule exocytosis, involving the release of perforin and GrB. It is known from influenza vaccine research that GrB correlates with protection and enhanced CTL response to influenza vaccination in older adults[60]. We looked at CTLs after day 7 of prime doses and found that combination immunization significantly increased differentiation of CTLs indicated by GrB+ expression (Figure7E-H) and perforin (Figure 7I-J) upregulation within one week after initial vaccine doses. Seven days after mice groups received booster doses that were administered on day 14, we found a robust 10-fold upregulation of GrB+ positive CD8+ T-cells indicating successful vaccine elicited differentiation of cytotoxic T-cells (Supplementary Figure 8).

Interestingly, it has been reported that cytotoxic CD4+ T-cells can compensate for age related decline of immune cell protection such as B cell loss and a less robust antibody response[61]. Strikingly, we found in SV.Spike+αOX40 immunized mice showed a significant increase of cytotoxic CD4+ T-cells indicating that our vaccine not only induced Th1-type CD4+ T helper functions but has the potential to improve direct CD4+ T-cell mediated virus-killing, thus, adding an extra layer to immune protection against SARS-CoV-2 in more vulnerable older populations. One important early feature of response to the SV.Spike+αOX40 immunization is a strong interferon-gamma (IFNg) secretion (Figure 7K), which is associated with polarization to Th1-type effector cells and cytotoxic T-cells. In order to investigate the recruitment and specificity in CTLs to prevent SARS-CoV-2 cell entry, we analyzed the potential of T-cells isolated from SV.Spike and/or αOX40 immunized and naïve mice on day 7 after prime doses to block the infection of 293T cells with SARS-CoV-2-spike expressing, luciferase-encoding pseudovirus. VSVG expressing, luciferase-encoding pseudovirus was used as control. We found that splenic T-cells from SV.Spike and SV.Spike+αOX40 mice potently inhibited infection with SARS-CoV-2 pseudotyped lentivirus (Figure 7L) compared to control (Figure 7M). In conclusion, SV.Spike+αOX40 activated T-cells display a Th-1 effector phenotype that promotes effector differentiation and direct T-cell mediated cytotoxicity against SARS-CoV-2 spike within one week after prime vaccine doses.

### 2.8 SV.Spike in combination with αOX40 drives metabolic activation of B cells and T-cell dependent B cell support

Almost all durable neutralizing antibody responses as well as affinity matured B cell memory depend on CD4+ T-cell helper. GSEA of RNAseq data between T-cells from the SV.Spike+αOX40 vaccinated and naive group one week after prime vaccine doses revealed selective enrichment of the gene set characteristic for activation of B cells (Figure 8A) (p<0.05). To test if SV.Spike combination with αOX40 selectively regulates T-cell dependent B cell activation, we investigated CD4+ T-cell activation and differentiation in mice vaccinated with SV.Spike and/or αOX40 one week after booster vaccine doses by flow cytometry analysis. We found that SV.Spike+αOX40 immunized mice had a 3-fold significant increase of overall CD44+positive splenic CD4+ T-cells compared to naïve mice (Supplementary Figure 9). We next analyzed follicular CD4+ T helper (Tfh) cells that are a subset of CD4+ T-cells required for most IgG responses promoting high-quality neutralizing antibodies and we found a 3-fold increase of ICOS+CXCR5+ (Figure 8B, C) and a 2 fold increase CD44+CXCR5+ (Figure 8D, E) positive CD4+ T-cells in splenocytes from the SV.Spike+αOX40 group indicating Tfh cell differentiation. We isolated B cells from spleens and performed a metabolic flux analysis on day 21 after initial vaccine doses and we found that isolated B cells from SV.Spike+αOX40 immunized mice were metabolically reprogrammed indicating potent vaccine elicited B cell activation. Activated B cells in the combination immunized group experienced a 2.5-fold increase in mitochondrial respiration (Figure 8F, G) and glycolysis (Figure 8G, H) when compared to B cells isolated from mice spleens that were vaccinated with a single agent or compared to naïve mice. Association analysis of the frequencies of Tfh cells with SARS-COV-2 spike IgG antibody titers revealed that Tfh cells positively correlated with the SARS-CoV-2 spike IgG serum levels in the SV.Spike (R^2^ = 0.9722, P=0.002) and SV.Spike+αOX40 group (R^2^ = 0.83, P = 0.0290) with the highest amounts of IgG antibodies and Tfh cells in the combination (Figure 8I). Taken together, these results indicate SV.Spike+αOX40 vaccine induced the most potent T-cell dependent B cell response.

**Figure 8.**
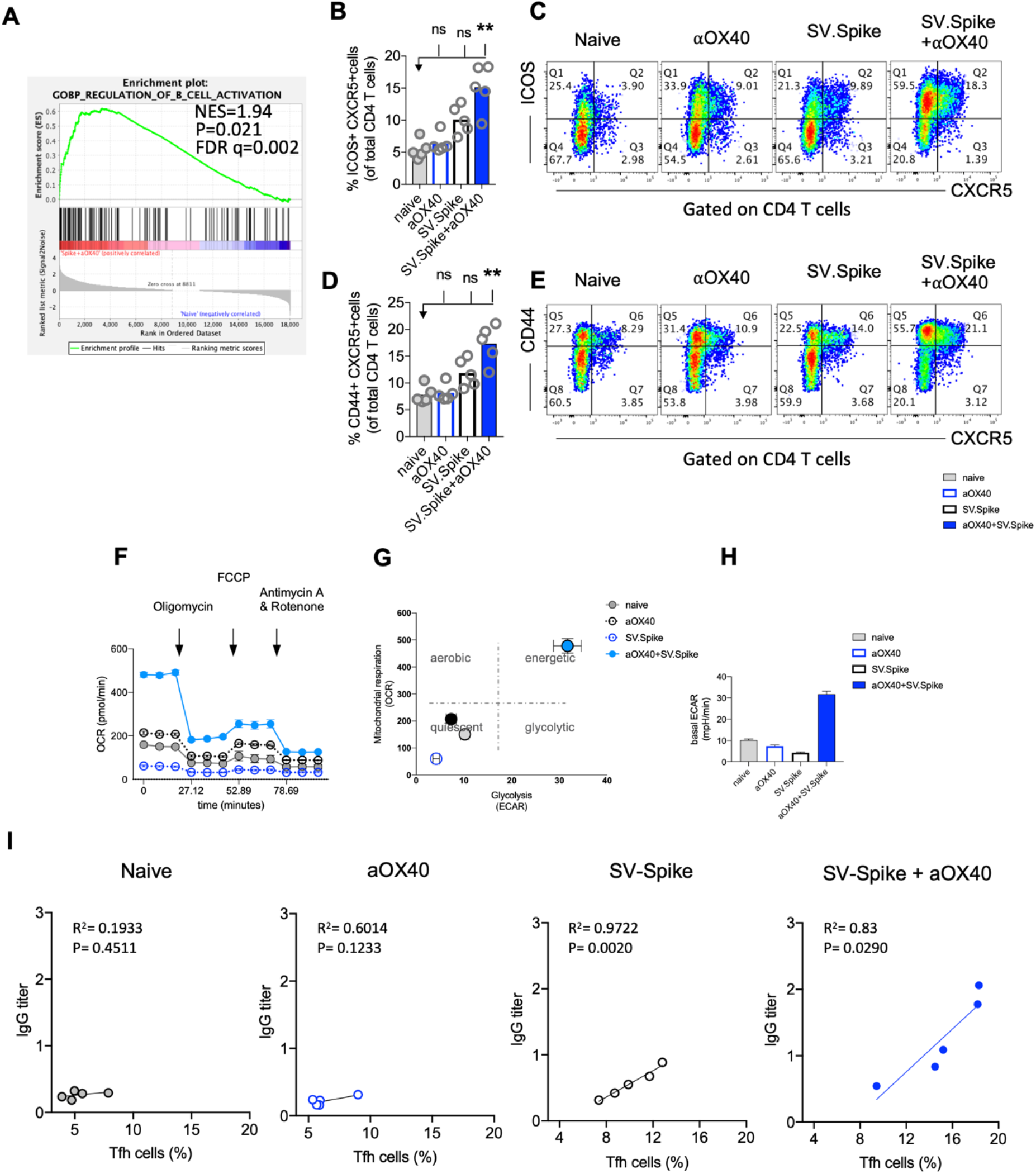
SV.Spike in combination with αOX40 drives follicular T helper cell function and metabolic activation of B cells. C57BL/6J mice were vaccinated with SV.Spike and/or αOX40. Naive mice were used as control. T-cells were isolated on day 7 after prime vaccine doses and RNAseq was performed **(A)**. GSEA for biological processes identified pathway enrichment that regulates B cell activation after prime vaccine doses in combination immunized mice. Splenocytes were excised on day 21 for flow cytometry analysis **(B-E)**. **(B-C)** CXCR5+ICOS+ expressing CD4+ T-cells and **(D-E)** CXCR5+CD44+ expressing CD4+ T-cells indicating Tfh-cell differentiation with representative plots (n=5 individual mice per group). **(F-H)** B cells were isolated for Seahorse metabolic flux analysis one week after boost doses. **(F)** Mitochondrial respiration was assessed by measuring the median values of oxygen consumption rates (OCR) in B cells of indicated groups using an extracellular flux analyzer. Oligomycin, FCCP, Antimycin A and Rotenone were injected as indicated to identify energetic mitochondrial phenotypes. **(G)** Energy Map (OCR versus ECAR) of B cells from naïve or mice treated with SV.Spike and/or αOX40 on day 21. **(H)** Baseline extracellular acidification rates (ECAR) in B cells of indicated groups. Error bars indicate SEM. Results are representatives of one or two independent experiments. Bars or symbols represent means ± SEM, and statistical significance was determined with the Kruskal-Wallis test followed by the he Dunns’ test. n.s. > 0.05, **p<0.005. **(I)** Correlation analysis of ICOS+CXCR5+ expressing Tfh cells with IgG antibody titers at 21 days post vaccination. (n=5). Pearson’s rank correlation coefficients (R) and p values are shown.

### 2.9 Combination of SV.Spike and αOX40 promotes robust T-cell specific immune response in lungs

Most vaccines for airborne infectious diseases are designed for delivery via the muscle or skin for enhanced protection in the lung. We investigated if SV.Spike vaccine-induced T-cells can readily home most efficiently to the lungs prior to and shortly after pathogen exposure. To address the immune responses in the lungs, we immunized mice with SV.Spike and/or αOX40 and excised PBS-perfused lungs one week after booster doses for single cell suspensions and performed flow cytometry staining (Figure 9, Supplementary Figure 9). We found an increase of ICOS+ CXCR5+ double-positive T helper cells indicating presence of B cell supporting Tfh cells in the SV.Spike single agent and combination immunized group. We further found an increase of Th-1 type effector CD4+ T-cells in lungs from combination treated mice indicated by expression of ICOS+Tbet+ double-positive effector CD4+ T-cells (Figure 9C, D). We next investigated if effector CTLs were successfully recruited into the lungs after 3 weeks of initial vaccine administration. While we found the highest increase of differentiated cytotoxic CD4+ T and CD8+ T-cells in lungs from the combination treated group (Figure 9E-H, Supplementary Figure 8), we observed a significant increase of differentiated cytotoxic CD8+ T-cells homing in the lungs of the SV.Spike single agent immunized group, although this increase was less pronounced compared to the combination group. These data indicate a successful recruitment of vaccine mediated antiviral Th1-type effector T-cells to the lungs.

**Figure 9.**
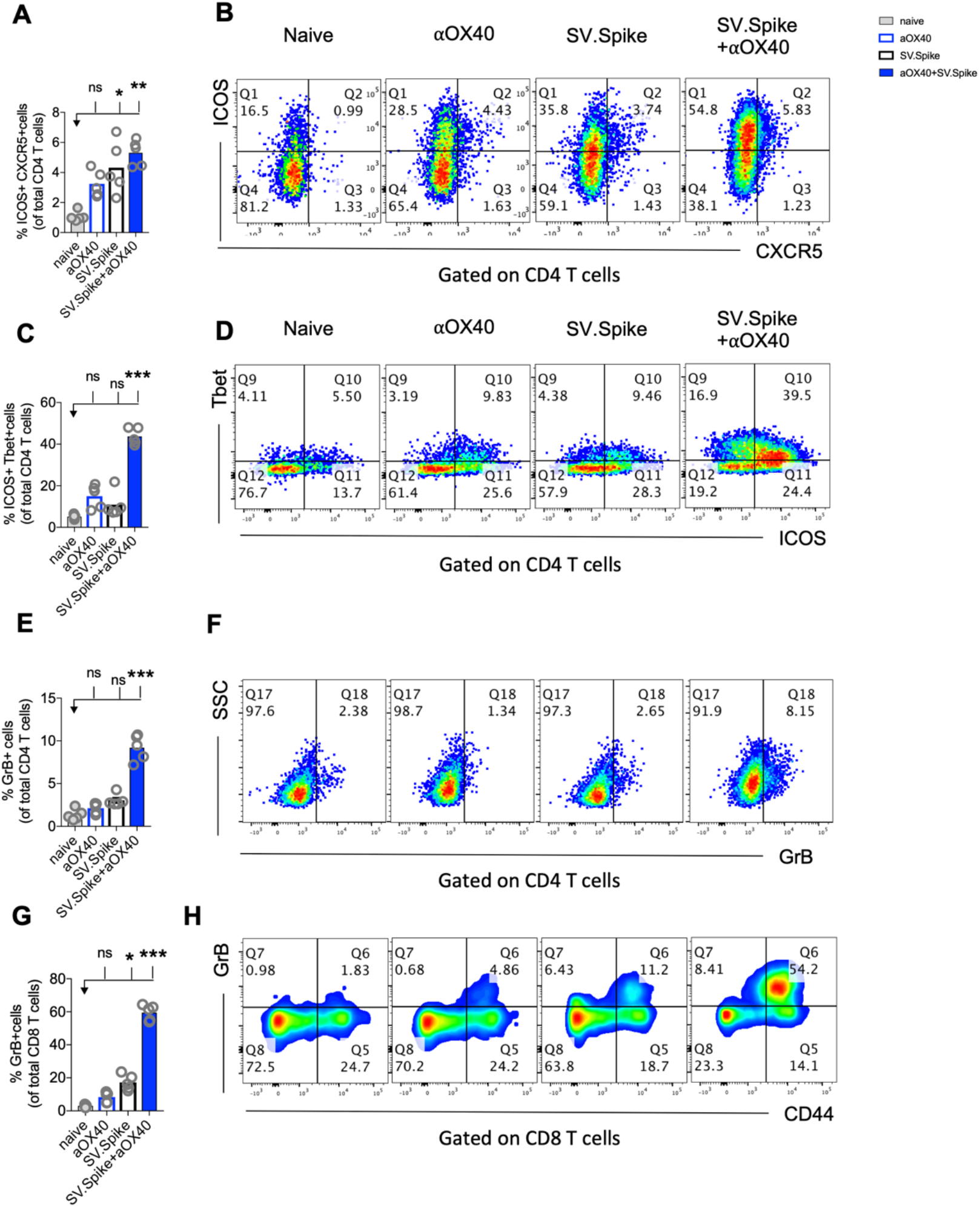
Combination of SV.Spike and αOX40 promotes robust tissue specific Th1-type T-cell immune response in lungs. Presence of activated T-cells in lungs after 21 days after prime vaccine doses indicate tissue specific immune protection. C57BL/6J mice were immunized by a Prime/Boost strategy with SV.Spike and/or αOX40 and lungs were excised and a single cell-suspension was stained for flow cytometry analysis. Naive mice were used as control. **(A)** CD4+ Tfh type T-cells presence in the lung indicated by ICOS+CXCR5+ double-positive CD4+ T-cells. **(B)** Representative plots. **(C)** Expression of ICOS+Tbet+ double positive CD4+ T-cells indicating Th-1 type effector cells polarization and recruitment to the lungs. **(D)** Representative plots. (E-H) Cytotoxic T-cells in lungs indicated by **(E)** Granzyme B positive CD4+ T-cells and representative plots (F) and CD8+ effector T-cells indicated by GrB+ and representative plots **(G, H)**. Bars or symbols represent means ± SEM. Each symbol represents one individual mouse. Statistical significance was determined with the Kruskal-Wallis test followed by the he Dunns’ test. n.s. > 0.05, *p<0.05, **p<0.005, ***p≤ 0.001.

### 2.10 SV.Spike and αOX40 promotes CD4+ T-cell memory formation and long-term protection upon re-challenge with SARS-CoV-2 spike antigen

Boosting both, local and systemic memory T-cell response is a useful strategy to achieve long term immunity. We analyzed development of T-cell memory in spleens fourteen weeks after initial prime vaccine doses of SV.Spike and/or αOX40 prime-boost immunized mice by flow cytometry. We found that mice in the SV.Spike+αOX40 combination group developed significant effector CD4+ T memory indicated by CD44+ CD62L+ double-positive CD4+ T-cells (Figure 10 A-C) compared with naïve mice, reiterating the importance of the combination vaccination in generating strong immune responses memory protection from infection and/or disease against SARS-CoV-2.

**Figure 10.**
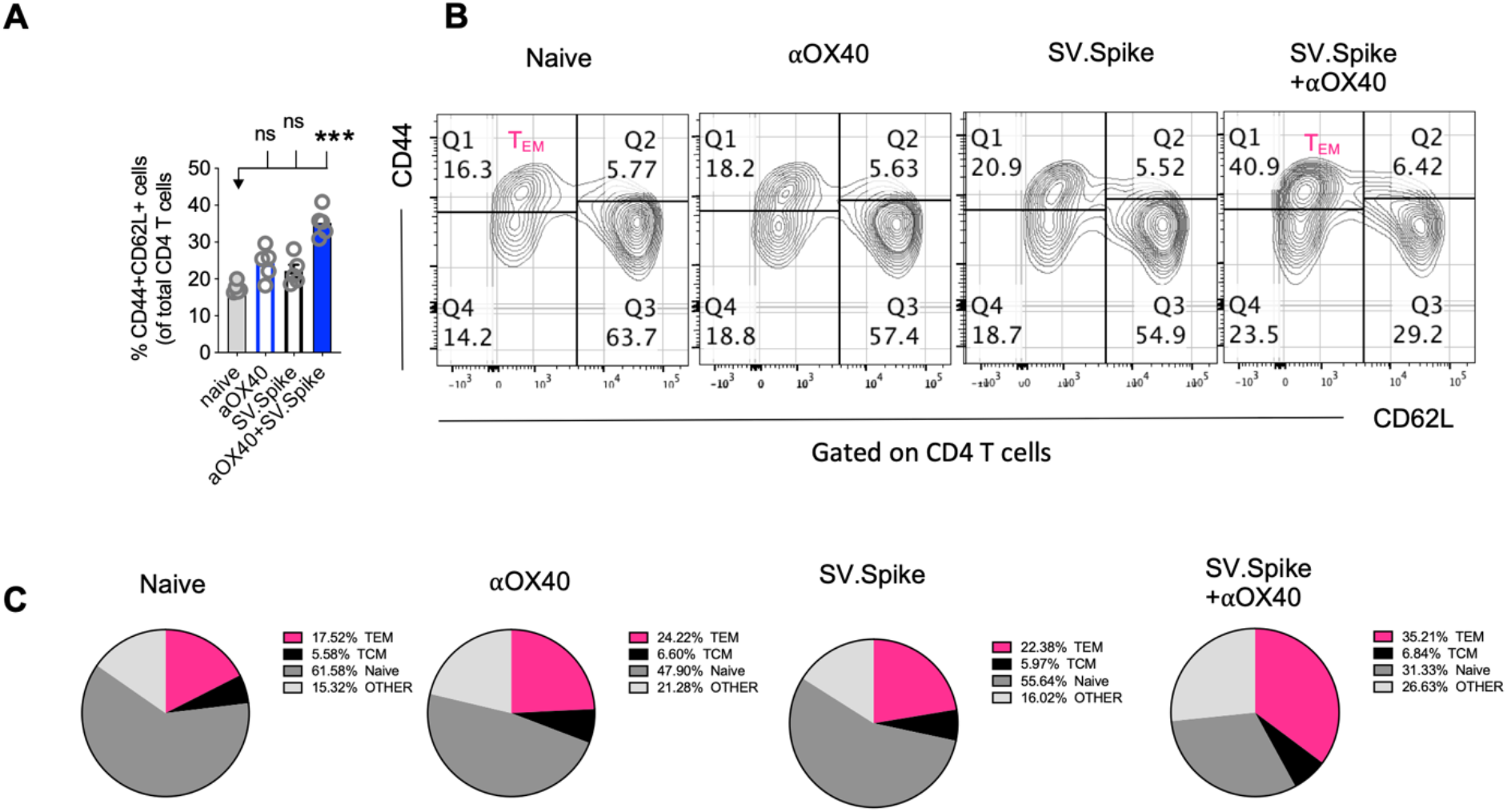
Combination of SV.Spike and αOX40 potentiates CD4 effector memory T-cells 14 weeks after prime vaccine doses. Splenocytes from indicated immunized C57BL/6J mice groups were harvested 14 weeks after first vaccine doses. Memory phenotype was characterized in spleen from indicated groups by flow cytometry by gating on CD4+ cells. The percentage of CD4+ T-cells expressing CD62L and/or CD44 was analyzed and shown **(A)**. **(B)** Representative contour plots and **(C)** pie charts. (n=5 mice per group). TCM, central-memory T-cells; TEM, effector-memory T-cells.

To further explore the long-term protection efficacy of our SV.Spike vaccine against SARS-CoV-2 virus challenge, C57BL/6J mice (n = 5 each group) received prime and boost immunizations of SV.Spike and/or αOX40 and placebo (naïve group) via the i.p. route. At day 100 post-immunization, we additionally administered one dose of SV.Spike, to recapitulate Spike antigen endogenous entry through SV vector injection (Figure 11A). Spleens or sera from re-challenged mice were collected 3 days after SARS-CoV-2 spike antigen injection and processed for T-cell response analysis (Figure 11B-F, Supplementary Figure 10) and detection of specific anti-spike protein IgA, Ig and IgG isotypes by ELISA (Figure 11G). The SARS-CoV-2 pseudotyped lentivirus infectivity assay revealed that mice immunized with SV.Spike or SV.Spike and αOX40 are effective in reactivating circulating cytotoxic T-cells (CTLs) upon challenge with Spike antigen (Figure 11B). CTLs reactivation was also observed by flow cytometry as indicated by granzyme B upregulation in mice receiving combination vaccination (Figure 11C, D). Moreover, immunophenotyping analysis showed that CXCR5-ICOS-double-positive Th1-type effector CD4+ T-cells were strongly rebooted in re-challenged mice receiving SV.Spike combination vaccination compared to the same group of unchallenged mice (Figure 11E, F).

**Figure 11.**
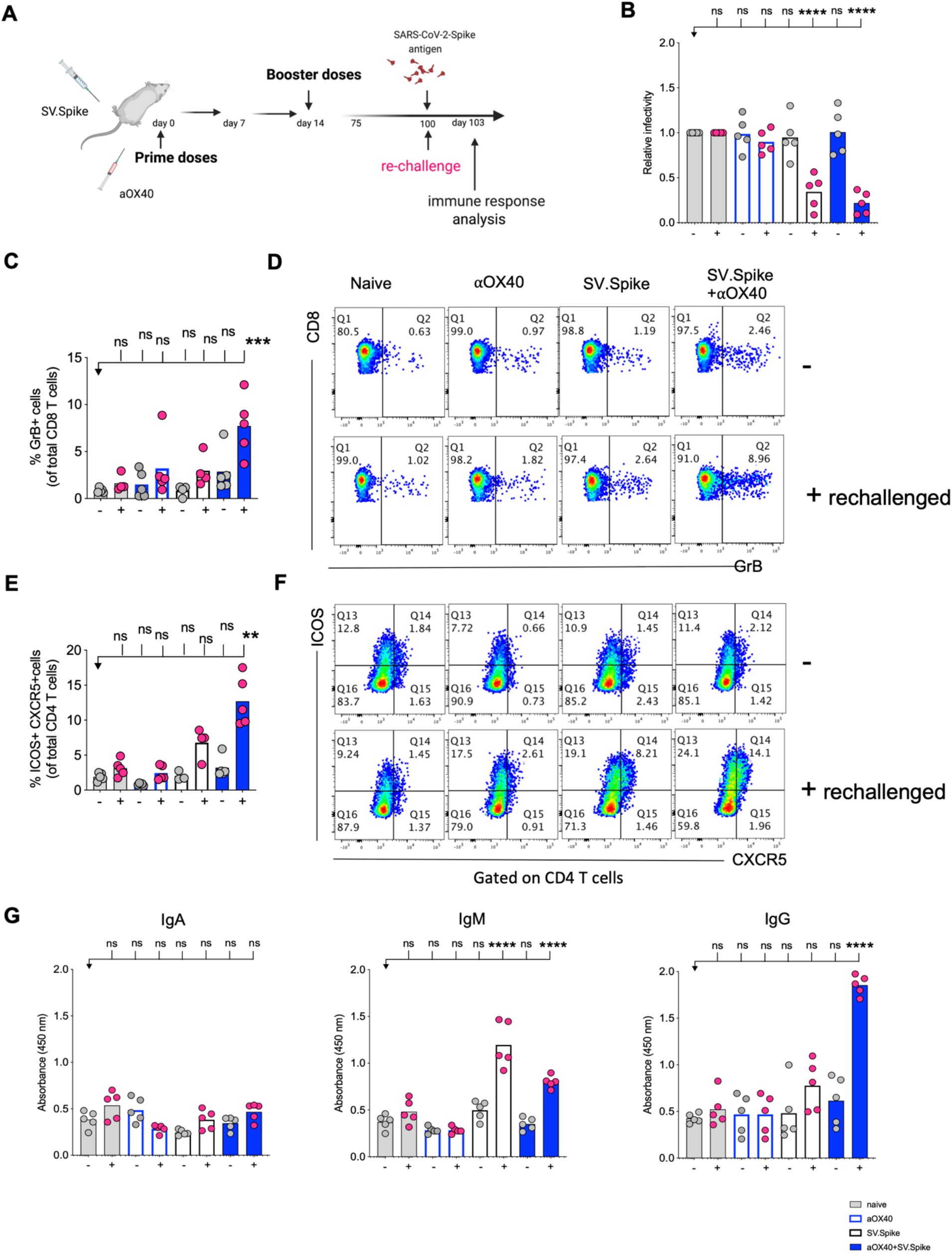
Challenging immunized mice with spike antigen promotes a fast and coordinated response of the two arms of the adaptive immune system. Humoral and T-cell immune responses were assessed in vaccinated mice after rechallenge with Sindbis carrying SARS-CoV-2-spike (SV.Spike). **(A)** Design steps of the rechallenge experiment in vaccinated of immunized C57BL/6J mice evaluated by **(B)** T-cell cytotoxic assay, **(C-F)** Flow cytometry indicating cytotoxic CD8 T-cell effector response by GrB+ positive CD8 T-cells and activation of CXCR5+ICOS+ positive Tfh cells upon rechallenge, **(G)** binding IgA, IgM, IgG antibody ELISA to SARS-CoV-2-spike recombinant protein (n=5 mice per group, or as otherwise indicated). Each symbol represents one individual mouse. Bars or symbols represent means ± SEM, and statistical significance was determined with one-way ANOVA with the Bonferroni correction **(B, G)** or with the Kruskal-Wallis test followed by the he Dunns’ test **(C-F)**. n.s. > 0.05, **p<0.005, ***p≤ 0.001, ****p ≤ 0.0001.

Antibody response analysis showed that immunization with SV.Spike or SV.Spike+αOX40 followed by Spike antigen injection induced strong production of IgM antibodies compared to the mice which did not received the antigen and the Naïve groups, and that was particularly evident in mice vaccinated with SV.Spike (Figure 11G). Strikingly, we noticed that combination of SV.Spike and αOX40 followed by challenge with antigen stimulated a high peak of Spike-specific IgG antibodies levels, which were about 4 times higher than the IgG levels of unchallenged mice and control group. No significant difference in the Spike-specific IgG response was detected in SV.Spike or single αOX40 re-challenged mice compared to the respective unchallenged mice and the control groups, whereas no SARS-CoV-2 spike-specific IgA were not detected in any of the groups (Figure 11G). Together, these data suggest that combination vaccination with SV.Spike and αOX40 conferred remarkably long-term and specific protection against SARS-CoV-2 infection by eliciting a durable humoral and T-cell response.

## 3 Discussion

The COVID-19 pandemic has placed substantial pressure on health systems to deliver an effective, and scalable vaccine that can be produce in hundreds of millions of doses. New vaccine platforms, reverse genetics, computational biology, protein engineering and gene synthesis facilitated this effort with successful production of several vaccines with that met these goals[62]. Over 162 candidates are undergoing preclinical development of which 53 already in clinical development (WHO https://www.who.int/publications/m/item/draft-landscape-of-covid-19-candidate-vaccines) and several have been administered to significant, if vastly incomplete, number of people. The latter include vaccine platforms based on DNA or RNA (Moderna[43], CureVac, BioNTech/Pfizer[63] adenovirus vector-based vaccines (CanSinoBIO[64], University of Oxford/AstraZeneca[65], Janssen Pharmaceutical Companies), inactivated vaccines (Sinopharm and Sinovac, Wuhan Institute of Biological Products), and protein subunit vaccines (Sanofi Pasteur/GSK, Novavax[66], Clover Biopharmaceuticals/GSK/Dynavax).

Despite promising results of early clinical trials of several vaccine candidates against SARS-CoV-2, there are still concerns regarding both safety and durability of the immune responses. Consequently, it is necessary to develop additional and improved vaccine candidates. An ideal vaccine against SARS-CoV-2 would be effective after one or two immunizations, conferring long-term protection to target populations such as the elderly or immunocompromised individuals, and reducing onward transmission of the virus to contacts[65]. It would protect against a broad range of coronaviruses and evolving variants, i.e., offer pancoronavirus protection. The benefit of developing such a vaccine would be even greater if it were available to be rapidly deployed in time to prevent repeated or continuous epidemics, economical and readily distributable worldwide without temperature constraints that limit access. This supports the use of alphavirus vaccine platforms that are rapid and straightforward to produce inexpensively, with less challenging temperature requirements, and with previously proven safety and efficacy.

The alphavirus-based replicon platform technology has been developed as vaccine candidates for many different infectious diseases, including influenza A virus (IAV), respiratory syncytial virus (RSV)[67; 68] Ebola (EBOV), hepatitis C virus (HCV), chikungunya (CHIKV, now in phase III)[69; 70] HIV (now in phase I), human papilloma virus (HPV, now in therapeutic phase II)[71]. Given the generic design of these platform and that new constructs can be made rapidly with synthetic design of the insert, it can be readily adapted to SARS-CoV-2 as we have demonstrated here. Moreover, when new virus species emerge, a vaccine platform that can be rapidly adapted to emerging viruses is highly desirable.

Sindbis virus and other alphaviruses have a natural tropism for lymphatic tissues and dendritic cells, relative resistance to interferon, high expression levels, lack of pre-existing anti-vector immunity in most human and animal populations, and efficient production of methodology in cell lines, with an accepted regulatory pedigree[72]. These observations indicate that a vaccine platform based on Sindbis virus vectors could contribute significantly to dealing with current and future vaccine needs. SV vectors constitute a novel alphavirus development platform that can be readily adapted to new pathogens and block emerging future pandemics early on in outbreaks. In nature SV has the safest profile among alphaviruses. SV is an RNA virus without replicative DNA intermediates and poses no risk of chromosomal integration or insertional mutagenesis. Hence, its presence is transitory. To avoid even transient adverse effects, our vectors have been attenuated by splitting the SV genome and by removing the packaging signal from the genomic strand that encodes the structural genes. Moreover, the combination of SV vectors with immunomodulatory antibodies like αOX40 makes them extremely effective.

Neutralizing antibodies (NAbs) have conventionally been the desired outcome of vaccination, as they are capable of intercepting and neutralizing microbes and their components as well as eliciting destructive anti-microbial innate immune responses[73]. Nonetheless, humoral immunity can decline over time and, as seen with influenza, can only last as short as one season. Many newer vaccines and vaccines in development are also designed to generate T-cell responses that have the potential to help the antibody response, promote long-term immune memory, have direct effector functions themselves, or activate innate effector cells such as macrophages and neutrophils[45; 74].

Here, we developed a Sindbis-based Spike-encoding RNA vaccine against SARS-CoV-2 and demonstrated that immunization with SV vector expressing SARS-CoV-2 spike along with a costimulatory agonistic αOX40 antibody induced a synergistic T-cell and antibody response and provided complete protection against authentic SARS-CoV-2 challenge in hACE2 transgenic mice. Our adaptable approach has the potential to boost tissue specific immunity and immune memory against respiratory viruses and aims to develop vaccines that could protect for several seasons or years. As a viral vector, we found that a Sindbis vector expressing SARS-CoV-2 spike antigen in combination with αOX40 markedly improves the initial T-cell priming, compared with the viral vector alone, which results in a robust CD4+ and CD8+ T-cell response and stable SARS-CoV-2 specific neutralizing antibodies. The vaccine efficiently elicits effector T-cell memory in respiratory tissues with a potential for long lasting protection against COVID19, which might extend for several years, through multiple beneficial mechanisms. It protects against infection with authentic, live SARS-CoV-2 preventing morbidity and mortality.

It has been shown that αOX40 controls survival of primed CD8+ T-cells and confers CTL-mediated protection[31; 75]. CTLs are a critical component of the adaptive immune response but during aging, uncoordinated adaptive responses have been identified as potential risk factors that are linked to disease severity for the outcome of COVID19 patients. It is known from influenza vaccine research that Granzyme B correlates with protection and enhanced CTL response to influenza vaccination in older adults. We looked at cytotoxic T-cells (CTLs) and found that combination vaccination significantly increased CD8+ cytotoxic T-cells indicated by granzyme B and perforin upregulation. Almost all durable neutralizing antibody responses as well as affinity matured B cell memory depend on CD4+ T helper cells. We found in combination vaccinated mice a significant increase of cytotoxic CD4+ T-cells indicating that our vaccine not only induced CD4+ T helper functions but has the potential to improve direct CD4+ T mediated virus-killing adding an extra layer to long-term immunity/protection in more vulnerable older populations.

Virus-specific CTL are quickly recruited to influenza-infected lungs by a Th1 response, specifically due to the production of IFNg[59]. IFNg regulates various immune responses that are critical for vaccine-induced protection and has been well studied[76; 77]. In a clinical trial of the now approved BNT162b1 IFNg secreting T-cells increased in participants 7 days after boost [45]. In this regard, one important early feature of the response to the SV.Spike+αOX40 immunization is a strong interferon-gamma (IFNg) secretion. We found a significant increase of CXCR3 and CX3CR1 positive expressing CD4+ T-cells, indicating effective recruitment and mobility of generated effector Th1 type T-cells in mice. This recruitment positively correlates with vaccine induced long-term immune protection and generation of neutralizing antibodies against SARS-CoV-2.

Both humoral and cell-mediated immune responses have been associated with vaccine-induced protection against challenge or subsequent re-challenge after live SARS-CoV-2 infection in recent rhesus macaque studies [78; 79] and there is mounting evidence that T-cell responses play an important role in COVID-19 mitigation[3; 80; 81]. We demonstrated that two doses of SV.Spike with or without αOX40 candidate vaccines induced neutralizing antibody titers in all immunized mice, with a strong IgG response in the mice receiving combination vaccination. Moreover, our data show that SV.Spike+αOX40 skewed Tfh cells toward CXCR5^+^ Tfh differentiation, which positively correlated with the magnitude of IgG isotype response. These findings indicate that the induction of CXCR5^+^ Tfh cell differentiation through vaccination may be beneficial for eliciting broad and specific NAb responses. Importantly, the synergistic activity of combination vaccination elicited antibodies that were able to efficiently neutralize SARS-CoV-2 pseudotyped lentivirus in all the mice tested. In addition, we show SV-Spike-based re-challenge in mice immunized with combination vaccination led to enhanced cytotoxic reactivation of T-cells and increased IgG seroconversion and response, and provided protection against re-challenge, reiterating the importance of the involvement of both humoral and cellular immune responses in SARS-CoV-2-mediated immunity.

The SV.Spike platform evaluated in this study has the advantage that it is inexpensive, stable, easy to produce. Cost projections based on using our upstream and downstream processes for production of a SV based vaccine are in line with or below costs per dose for other vaccines in use today. Moreover, unlike other mRNA vaccine candidates this viral platform does not require a cold-chain during transportation and storage. It can be easily reconstitute after lyophilization process and is suitable for rapid adaptation such that potential new viruses/threats in an emerging outbreak can be rapidly targeted[82]. Thus, for emerging pathogens like SARS-CoV-2, the SV platform can be an efficient and cost-effective alternative to the traditional large-scale antigen production or technology platforms that require extended time for implementation. Development of a successful SV vector vaccine is readily translatable into human vaccination efforts.

As shown in this study, SV.Spike can be applied alone or can be combined with immunomodulatory reagents like αOX40 in a remarkably efficient prime-boost regimen. Our goal is to exploit the combined SV.Spike + αOX40 formulation and integrate the two components into a single vector, to further facilitate administration and immunomodulatory response. Our lab has recently demonstrated that the expression of full-length antibodies from SV vectors is feasible and effective and that we can also integrate a third gene of interest such as an antigen or a cytokine (unpublished). Taken together, these data provide an insight into antigen design and preclinical evaluation of immunogenicity of SV-based vaccines, and support further development of SV.Spike as a vaccine candidate for protection against COVID-19 and further to generate a pancoronavirus vaccine.

## 4 Material and Methods

### 4.1 Cell lines

Baby hamster kidney (BHK) and 293T-cell lines were obtained from the American Type Culture Collection (ATCC). 293T/ACE2 cell line was obtained from BEI Resources.

BHK cells were maintained in minimum essential α-modified media (α-MEM) (Corning CellGro) with 5% fetal bovine serum (FCS, Gibco) and 100 mg/ml penicillin-streptomycin (Corning CellGro). 293T and 293T/ACE2 cells were maintained in Dulbecco’s modified Eagles medium containing 4.5 g/l Glucose (DMEM, Corning CellGro) supplemented with 10% FCS, 100 mg/ml penicillin-streptomycin. All cell lines were cultured at 37 °C and 5% CO2.

### 4.2 SV Production

SV.Spike expressing vector was produced as previously described[38; 39; 83; 84]. Briefly, plasmids carrying the replicon (pT7-SV-Spike) orT7-DMHelper RNAs were linearized with XhoI. In vitro transcription was performed using the mMessage mMachine RNA transcription kit (Invitrogen Life Sciences). Helper and replicon RNAs were then electroporated into BHK cells and incubated at 37°C in αMEM supplemented with 10% FCS. After 12 hours, the media was replaced with OPTI-MEM (GIBCO-BRL) supplemented with CaCl2 (100 mg/l) and cells were incubated at 37°C. After 24 hours, the supernatant was collected, centrifuged to remove cellular debris, and frozen at −80°C. Vectors were titrated as previously described [85].

### 4.3 Pseudotyped Lentivirus Production

SARS CoV-2 pseudotyped lentiviruses were produced by transfecting the 293T cells with the pLenti-Puro vectors (Addgene) expressing Luciferase or β-Galactosidase, with pcDNa3.1 vector expressing SARS-CoV-2 spike (BEI repository) and the helper plasmid pSPAX2 (Addgene). The VSV-G and empty lentiviruses were produced by replacing pCDNA3.1-Spike with pcDNA3.1-VSV-G or pCDNA3.1 empty vector, respectively (Addgene). The transfections were carried out using the Polyethylenimine (PEI) method with the ratio at PEI:pLenti:pcNDA3.1-Spike:pSPAX2 = 14:2:2:1 or PEI:pLenti:pVSV-G/pcNDA3.1:pSPAX2 = 10:1:0.5:3. The virus-containing medium was harvested 72 hours after transfection and subsequently pre-cleaned by centrifugation (3,000 g) and a 0.45 μm filtration (Millipore). The virus-containing medium was concentrated by using a LentiX solution (TakaraBio) a 10:1 v/v ratio and centrifuged at the indicated RCF at 4 °C. After centrifugation, the supernatant was carefully removed and the tube was drained on the tissue paper for 3 minutes. Dulbecco’s modified Eagles medium containing 4.5 g/l Glucose (DMEM) was added to the semi-dried tube for re-suspension and then stored at −80 °C.

### 4.4 Detection of SARS-CoV-2 spike pseudotyped lentivirus infectivity

*Luciferase*- and *nLacZ*-encoding SARS CoV-2 Spike or VSV-G pseudotyped lentivirus titers were determined making serial dilutions of the vectors in DMEM and infect 293T/ACE2 cells pre-plated in 96-well culture plates (10^4^ cells/well) and 24h later, fresh media was added. For *Luciferase*-encoding pseudotype, cells were lysed 72h later using cell lysis buffer and lysates were transferred into fresh 96-well luminometer plates, where luciferase substrate was added (Thermo Fisher), and relative luciferase activity was determined (Supplementary Figure 4C). For *nLacZ*-encoding pseudotypes, cells were washed with PBS and stained for 16h at 37 °C with X-Gal Solution [1 mg/ml X-Gal in PBS (pH 7. 0) containing 20 mM potassium ferricyanide, 20 mM potassium ferrocyanide and 1mM MgCl2] (Supplementary Figure 4D). Vector titers refer to the number of infectious particles (transducing units per milliliter of supernatant [TU/mL] and were estimated as the last dilution having detectable reporter activity. Correct assembling of pseudotypes was assessed by western blot following standard protocol, to detect the expression of SARS-CoV-2-spike and p24 proteins. SARS-CoV-2 spike (BPS Bioscience) and p24 (Abcam) recombinant proteins were used as positive controls (Supplementary Figure 4A, B).

### 4.5 *In vivo* experiments

All experiments were performed in accordance with protocols approved by the Institutional Animal Care and Use Committee of New York University Grossman School of Medicine. Six to 12-week old female C57BL/6J albino mice (B6(Cg)-Tyr<c-2J>/J,Cat#000058) and Hemizygous (B6(Cg)-Tg(K18-ACE2)2Prlmn/J; Cat#034860) (hACE2-Tg) mice expressing the human ACE2 receptor or non-carrier controls were purchased from Jackson Laboratory.

### 4.6 ABSL3 experiments using SARS-CoV-2 Coronavirus

Three weeks after prime and boost vaccination doses, hACE2-Tg and non-carrier control mice were challenged with 10^4^ pfu particles of SARS-CoV-2 Coronavirus via the intranasal (i.n.) route (Figure 4F). We recorded daily the body weight of each mouse after infection for a total of 14 days. The New York University Grossman School of Medicine (NYUSOM) Animal Biosafety Level 3 (ABSL3) Facility, located on the third floor of the Alexandria Center for Life Science West Tower, is a 3,000 sq. ft. high-containment research facility under the responsibility of the Office of Science & Research and its Director of High-Containment Laboratories. It has been designed and it is operated in compliance with the guidelines of the Centers for Disease Control and Prevention (CDC) and the National Institutes of Health (NIH). All research and non-research operations are governed by institutional standard operating procedures (SOPs). As per those SOPs, all users undergo specific training and require medical and respiratory protection clearance. The facility and its SOPs are re-certified by an outside consultant on a yearly basis. The NYUSOM ABSL3 has also been registered with the Department of Health and Mental Hygiene of the city of New York since March 2017.

### 4.7. Mouse vaccination and serum collection

Mice were i.p. immunized with SV.Spike (10^7^ TU/ml) in a total volume of 500 μl was injected i.p. into the left side of the animal. The immunostimulatory αOX40 antibody (clone OX-86, BioXCell) was injected i.p. into the left side of the animal at a dose of 250 μg per injection. Mice were boosted once at 2 weeks. Sera were collected at 7 days post-2^nd^ vaccination and used to detect neutralizing activity.

Therapeutic efficacy of vaccines was monitored in two ways: vaccinated hACE2-Tg mice that were challenged with SARS-CoV-2 Coronavirus in BSL3 were tested for survival compared to their non immunized control group. Survival was monitored and recorded daily.

### 4.8 *In vivo* delivery of nLacZ-SARS-CoV-2 pseudotype and X-Gal histochemistry

Isoflurane-anesthetized 4-week-old young adult hACE2-Tg mice were dosed intranasally with a 70-μl volume of *nLacZ*-encoding lentiviral vector (titer 5.18×10^3^ TU/ml). Isoflurane anesthesia (2.5% isoflurane/1.5l oxygen per minute) and dosing of animals was carried out in a vented BSL-2 biological safety cabinet. For processing of mouse lungs for X-Gal staining of intact tissue, lungs were inflated through the trachea with OCT embedding as described previously[86]. Intact airways were submerged in 0.5% glutaraldehyde for 2 h at 4 °C, washed in PBS/1 mM MgCl2 and stained for 16h at 37 °C with X-Gal Solution [1 mg/ml X-Gal in PBS (pH 7. 0) containing 20 mM potassium ferricyanide, 20 mM potassium ferrocyanide and 1mM MgCl2].

### 4.9 Neutralization experiments

#### 4.9.1 SARS-CoV-2 spike-hACE2 blocking assay

To measure protective NAbs, COVID-19 convalescent plasma was diluted (1:10) and incubated with recombinant SARS-CoV-2 full-length Spike (BPS Bioscience) for 1 h at 37 °C prior to adding to an hACE2 pre-coated ELISA plates. The NAb levels were calculated based on their inhibition extents of Spike and hACE2 interactions according to the following equation: [(1-OD value of samples/OD value of negative control) × 100%]. A neutralizing antibody against SARS-CoV-2 spike (Bio Legend) was used as a positive control.

#### 4.9.2 SARS-CoV-2 spike pseudotyped lentivirus inhibition assay

Pseudotyped lentivirus inhibition assay was established to detect neutralizing activity of vaccinated mouse sera and inhibitory ability of antiviral agents against infection of SARS-CoV-2 spike pseudotyped lentivirus in target cells. Briefly, pseudotyped virus containing supernatants were respectively incubated with serially diluted mouse sera at 37 °C for 1h before adding to target cells pre-plated in 96-well culture plates (10^4^ cells/well). 24h later, fresh media was added and cells were lysed 72h later using cell lysis buffer. Lysates were transferred into fresh 96-well luminometer plates. Luciferase substrate was added (Promega), and relative luciferase activity was determined. The inhibition of SARS-COV-2 Spike pseudotype lentivirus was presented as % inhibition.

### 4.10 Cell-cell fusion assay

The establishment and detection of several cell–cell fusion assays are as previously described [47]. In brief, 293T/ACE2 cells were used as target cells. For preparing effector cells expressing SARS-CoV-2 spike, 293T cells were transiently co-transfected with pCDNA3.1-Spike and pMAX-GFP or with pMAX-GFP only as control, and applied onto 293T/ACE2 cells after 48 h. Effector and target cells were cocultured in DMEM plus 10% FBS for 6 h. After incubation, five fields were randomly selected in each well to count the number of fused and unfused cells under an inverted fluorescence microscope (Nikon Eclipse Ti-S).

### 4.11 Inhibition of SARS-CoV-2-spike-mediated cell-cell fusion

The inhibitory activity of neutralizing antibodies from immunized mice sera on a SARS-CoV-2-spike-mediated cell–cell fusion was assessed as previously described[49; 87].

Briefly, a total of 2 × 10^4^ target cells/well (293T/ACE2) were incubated for 5 h. Afterwards, medium was removed and 10^4^ effector cells/well (293T/Spike/GFP) were added in the presence of serum from C57BL/6J immunized mice at 1:100 dilution in medium at 37 °C for 2 h. The fusion rate was calculated by observing the fused and unfused cells using fluorescence microscopy.

### 4.12 Immunocytochemistry

Cell immunocytochemistry was performed as described previously[88]. Briefly, cells were fixed with 4% paraformaldehyde (PFA) for 20 min at room temperature and then the membrane was permeabilized with 0.1% (vol/vol) Triton X-100 (Fisher Scientific). Incubation with blocking solution (5% normal goat serum) was performed at room temperature for 45 min. Anti-mouse SARS-CoV-2-spike (GTX, 1:100) and anti-rabbit hACE2 (Thermo Fisher,1:100) were applied overnight at 4 °C followed by incubation of appropriate secondary antibodies conjugated with fluorophores. Confocal images were captured using the Zeiss LSM-800 system.

### 4.13 Flow cytometry

For flow cytometry analysis, spleens were harvested from mice and processed as previously described[39]. Extracted lungs were chopped in small pieces and incubated with a digestive mix containing RPMI with collagenase IV (50 μg/ml) and DNAseI (20 U/ml) for 30 min at 37 °C. Spleens and lungs were mashed through a 70-μm strainer before red blood cells were lysed using ammonium-chloride-potassium (ACK) lysis (Gibco). Cells were washed with PBS containing 1% FCS and surface receptors were stained using various antibodies. Fluorochrome-conjugated antibodies against mouse CD3, CD4, CD44, CD38, ICOS, OX40, CD62L, Perforin, Granzyme B and Tbet, CXCR5 were purchased from Biolegend. Fluorochrome-conjugated antibodies against mouse CD8a were purchased from BD Biosciences. Fluorochrome-conjugated antibodies against CXCR3 and Ki67 were purchased from Thermofisher. Stained cells were fixed with PBS containing 4% Formaldehyde. For intracellular staining, the forkhead box P3 (FOXP3) staining buffer set was used (eBioscience). Flow cytometry analysis was performed on a LSR II machine (BD Bioscience) and data were analyzed using FlowJo (Tree Star).

### 4.14 T and B cell isolation

Total T-cells were freshly isolated with the EasySep™ mouse T Cell Isolation Kit. Total B cells were freshly isolated with the EasySep™ mouse B Cell Isolation Kit. Isolation of T and B cells were performed according to the manufacturer’s protocols (Stemcell Technologies).

### 4.15 Enzyme-Linked Immunospot (ELISPOT)

Enzyme-linked immunospot was performed as previously described[39]. Mouse IFNγ ELISPOT was performed according to the manufacturer’s protocol (BD Bioscience). Freshly isolated (1 x 10^5^) T-cells were directly plated per well overnight in RPMI supplemented with 10% FCS. No *in vitro* activation step was included. As positive control, cells were stimulated with 5ng/ml PMA+1μg/ml Ionomycin.

### 4.16 *Ex Vivo* Cytotoxic Assay

T-cells (8 × 10^5^/mL) from C57BL/6J immunized splenocytes were co-cultured with 293T/ACE2 cells (2 × 10^4^/mL), previously infected with 3×10^5^ TU of SARS-CoV-2 Luc-SARS-CoV-2 spike pseudotyped lentivirus. Cells were co-cultured in a 24-well plate for 2 days in 1 mL of RPMI 1640 supplemented with 10% FCS, washed with PBS and lysed with 100 μL of M-PER mammalian protein extraction reagent (Thermo Fisher) per well. Cytotoxicity was assessed based on the viability of 293T/ACE2 cells, which was determined by measuring the luciferase activity in each well. Luciferase activity was measured by adding 100 μL of Steady-Glo reagent (Promega) to each cell lysate and measuring the luminescence using a GloMax portable luminometer (Promega).

### 4.17 Transcriptome analysis of T-cells

Total RNA was extracted from freshly isolated T-cells on day 7 of treatment from spleens using RNeasy Kit (Qiagen). For each group, 5 C57BL/6J mice were used for biological repeats. RNA-seq was done by NYUMC Genome Center. RNA quality and quantity were analyzed. RNAseq libraries were prepared and loaded on the automated Illumina NovaSeq 6000 Sequencing System (Illumina). 1x S1 100 Cycle Flow Cell v1.5, 30 automated stranded RNA-seq library prep polyA selection, per sample.

### 4.18 RNA-Seq data analysis

RNA-seq data were analyzed by sns rna-star pipeline (https://github.com/igordot/sns/blob/master/routes/rna-star.md). Sequencing reads were mapped to the reference genome (mm10) using the STAR aligner (v2.6.1d)[89]. Alignments were guided by a Gene Transfer Format (GTF) file. The mean read insert sizes and their standard deviations were calculated using Picard tools (v.2.18.20) (http://broadinstitute.github.io/picard). The read count tables were generated using subread (v1.6.3)[90], (normalized based on their library size factors using DEseq2[91], and differential expression analysis was performed. To compare the level of similarity among the samples and their replicates, we used principal-component. All the downstream statistical analyses and generating plots were performed in R environment (v4.0.3) (https://www.r-project.org/). The results of gene set enrichment analysis were generated by GSEA software[92; 93]. The network of Gene Ontology terms was generated by Enrichment Map in Cytoscape. Additional protein–protein functional associations used in this study for bar graphs were retrieved from STRING (http://www.string-db.org/, version 11)[94], a well-known public database on several collected associations between proteins from various organisms.

### 4.19 Measurement of Oxygen Consumption and Extracellular Acidification Rates of T and B cells

T and B cell metabolic output was measured by Seahorse technology as previously described[95]. Purified T-cells from C57BL/6J immunized or control mice were plated at 6×10^5^ cells/well in a Seahorse XF24 cell culture microplate. Oxygen consumption rate (OCR) and extracellular acidification rate (ECAR) were measured using an Agilent Seahorse XFe24 metabolic analyzer following the procedure recommended by the manufacturer (Agilent). For the mitochondrial stress test, 1) oligomycin (1 μM), 2) FCCP (1.5 μM) and 3) rotenone (100 nM) and antimycin A (1 μM) were injected sequentially through ports A, B and C.

### 4.20 Immunoblot analysis

Western blot was performed to detect SARS-CoV-2 spike protein in 293T cells infected with SV.Spike and in the generated pseudotyped lentivirus. Cells were lysed in M-PER^®^ Mammalian Protein Extraction Reagent (Thermo Fisher) according to the manufacturer’s protocol. Lysates were separated by SDS-PAGE on 4-12% Bio-Rad gels, transferred to polyvinylidene difluoride (PVDF) membranes, blocked in 5% milk in TBS buffer with 0.1% Tween-20 (TBST). Primary antibodies to SARS-CoV-2 spike (GTX, 1:1000) and p24 (Abcam, 1:1000) were added overnight at 4 °C. HRP-conjugated secondary antibodies were added in 5% milk in TBST for 1 h at room temperature. BioRad Imaging System was used for visualization.

### 4.21 Statistical analysis

Statistical analysis was performed using GraphPad Prism 7.0 as described in figure legends. All data are shown as mean ± SEM. Figures were prepared using GraphPad Prism 7, Adobe Photoshop and ImageJ Software. Treated groups were compared using a one-way analysis using Prism7 (GraphPad Software) to naïve mice. Differences with a P value of <0.05 were considered significant: *P<0.05; **P<0.005; ***P<0.001.

### 4.22 Data Availability Statement

The original contributions presented in the study are included in the article/Supplementary Material, further inquiries can be directed to the corresponding authors.

## 5 Acknowledgments

Funding was provided by NIH 5R44CA206606 and by an NYU Grossman School of Medicine Institutional COVID-19 research fund. We would like to thank the NYU High Throughput Biology Laboratory for Seahorse usage, training and assistance and Dr. Shohei Koide for his contribution in reading the manuscript and providing helpful suggestions.

## 6 Author contribution

A.S., S.O. and D.M. conceived the study. A.S., S.O., A.H., designed experiments. A.S., S.O., A.M., C.P., Z.L. performed mouse experiments and/or data analysis. M.G.N., S.A.T. and K.A.S performed BSL3 experiments with live coronavirus and related data analysis. A.S., S.O. and D.M. wrote the manuscript. All authors reviewed and edited the manuscript.

