## Supplemental Figures for "Combination of a Sindbis-SARS-CoV-2 spike vaccine and αOX40 antibody elicits protective immunity against SARS-CoV-2 induced disease and potentiates long-term SARS-CoV-2-specific humoral and T-cell immunity"

**Supplementary Figures**

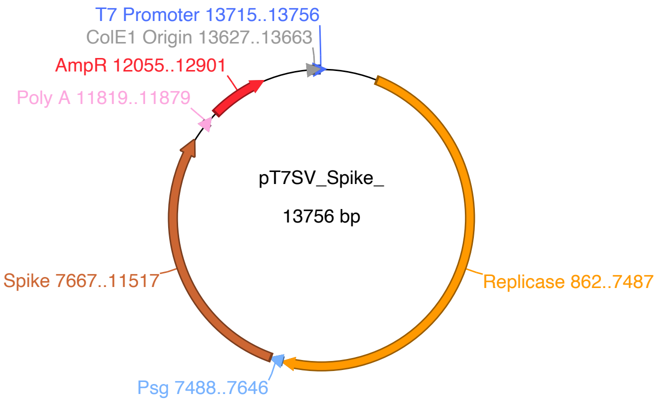


**Supplementary Figure 1.** SARS-CoV-2 spike sequences cloned into the SV vector expressing. The SARS CoV-2 spike sequence originates from the BEI Resource NR-52420 plasmid. The spike sequence was cloned into the *XbaI/ApaI* sites of the Sindbis replicon vector. The plasmid is linearized at the *XhoI* site, RNA is *in vitro* transcribed from the T7 promoter and capped. Electroporation of the replicon mRNA produces SV replicase that transcribes the spike gene from the subgenomic promoter (Psg).

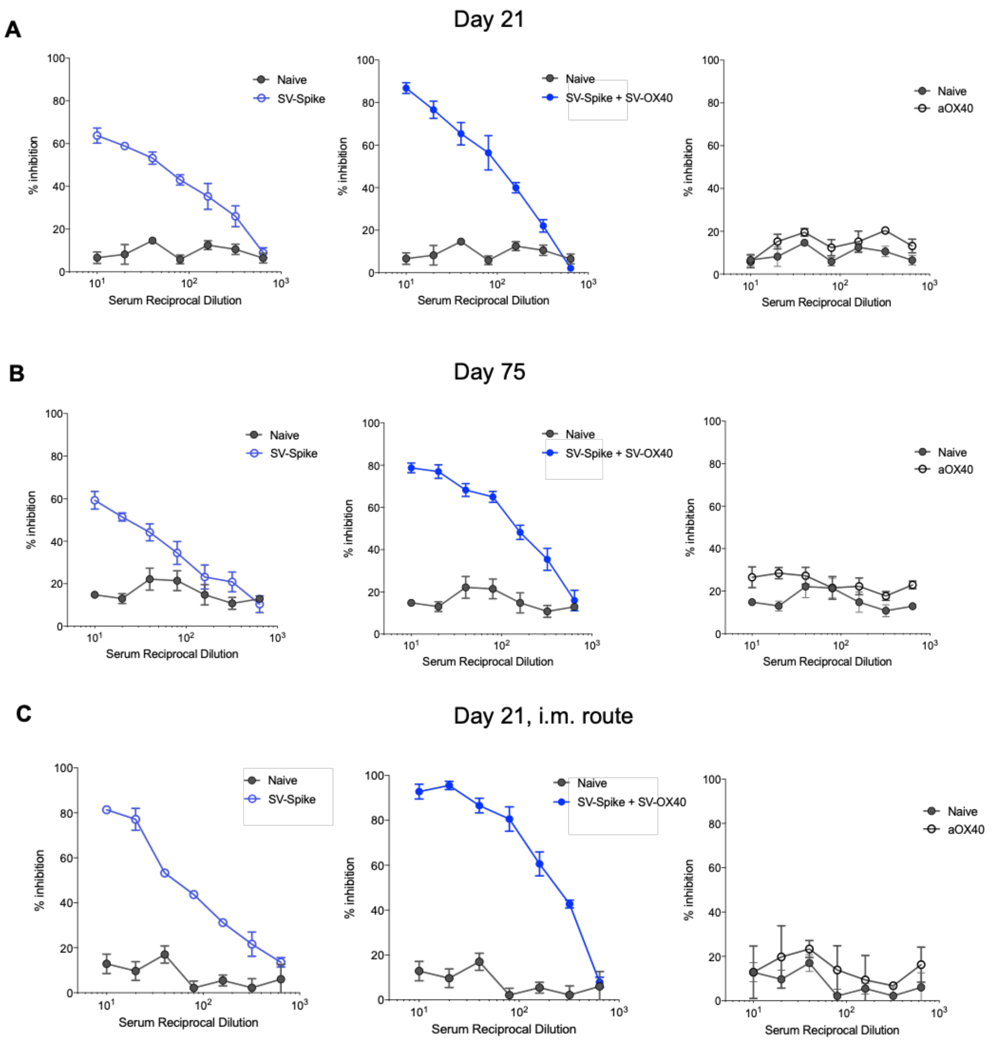


**Supplementary Figure 2.** Dose-response curves of anti-SARS-CoV-2 spike neutralizing antibodies blocking the SARS-CoV-2 spike-ACE2 binding. Inhibition curves of SARS-CoV-2 spike–hACE2 interaction for the indicated reciprocal serum dilutions by C57BL/6J mice sera collected at **(A)** 21 and **(B)** 75 days post vaccination with Sindbis expressing SARS-CoV-2 Spike (left panels) , SARS-CoV-2 Spike (middle panels) in combination with αOX40 and αOX40 alone (right panels) compared to the naïve group. Inhibition curves for mouse sera collected at 21 days post vaccination via the intramuscular (i.m.) route are shown in **(C)**. The data presented are the mean of n = 5 biological replicates with n = 2 technical replicates each curve.

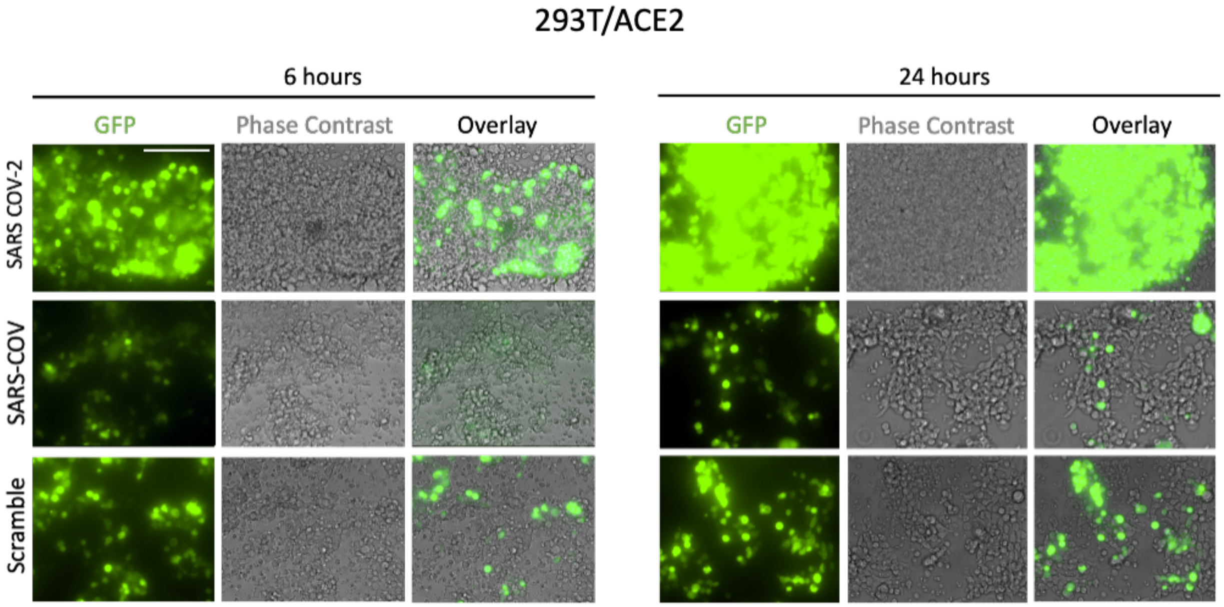


**Supplementary Figure 3**: Images of SARS-CoV and SARS-CoV-2 spike-mediated cell–cell fusion on 293T/ACE2 cells at 6 hours (left) and 24 hours (right). 293T have been co-transfected with pMAX-GFP/pCDNA3.1-SARS-COV or pMAX-GFP/pCAGGS-SARS-COV-2 Spike plasmids and applied onto 293T and 293T/ACE2 cells for the indicated time points. Scramble represents expression of GFP only. Scale bar, 100 µm.

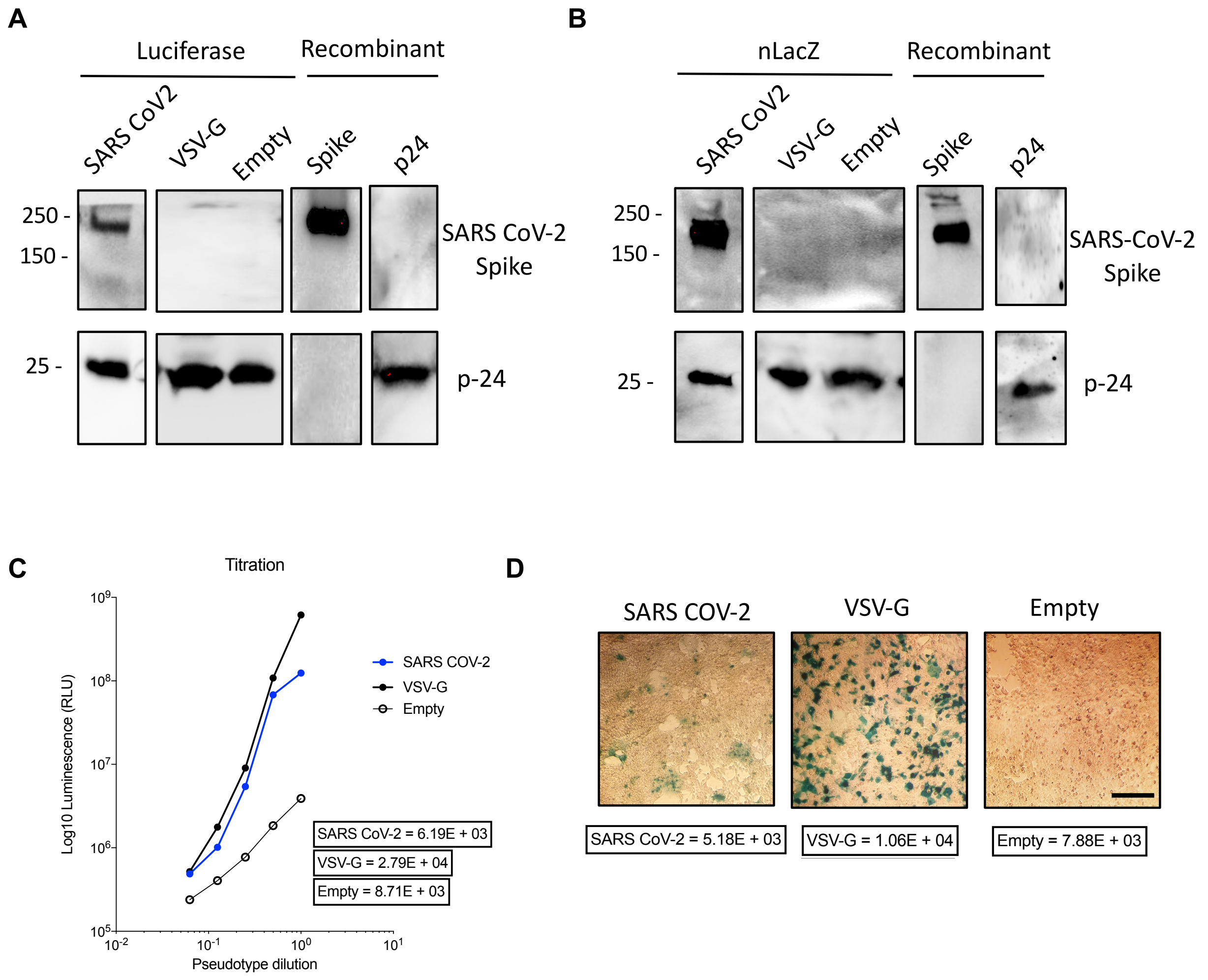


**Supplementary Figure 4:** Characterization of pseudotypes expressing SAR-CoV-2 Spike. Western blot analysis of the expression of p24 and Spike proteins from **(A)** Luciferase- and **(B)** nLacZ-encoding SARS-CoV-2 pseudotyped lentivirus produced in 293T cells transfected with pLentivirus expression plasmids as explained in the method section. A VSV-G encoding and empty (non-modified envelope) lentiviruses were also produced as expression controls. Purified SARS CoV-2 spike and p24 recombinant proteins were used as the positive controls. **(C)** Titration of Luciferase-encoding SARS-CoV-2 with spike protein, VSV-G and empty lentiviruses using HEK293 T/ACE2 cells. Log10 luminescence units (RLU) were measured. Titration values are expressed as TU/ml. n =3. **(D)** Immunofluorescence analysis of the expression of LacZ protein in HEK293 T/ACE2 cells mediated by SARS-CoV-2, VSV-G and empty pseudotyped particles. HEK293 T/ACE2 cells were infected with SARS-CoV-2-spike, VSV-G or empty pseudotype lentiviruses at 0.5 TCID50 per cell. Seventy-two hours later, cells were stained for X-Gal and observed microscopically. Cell nuclei were counterstained with Nuclear Fast Red Solution. Scale bar = 20 μm.

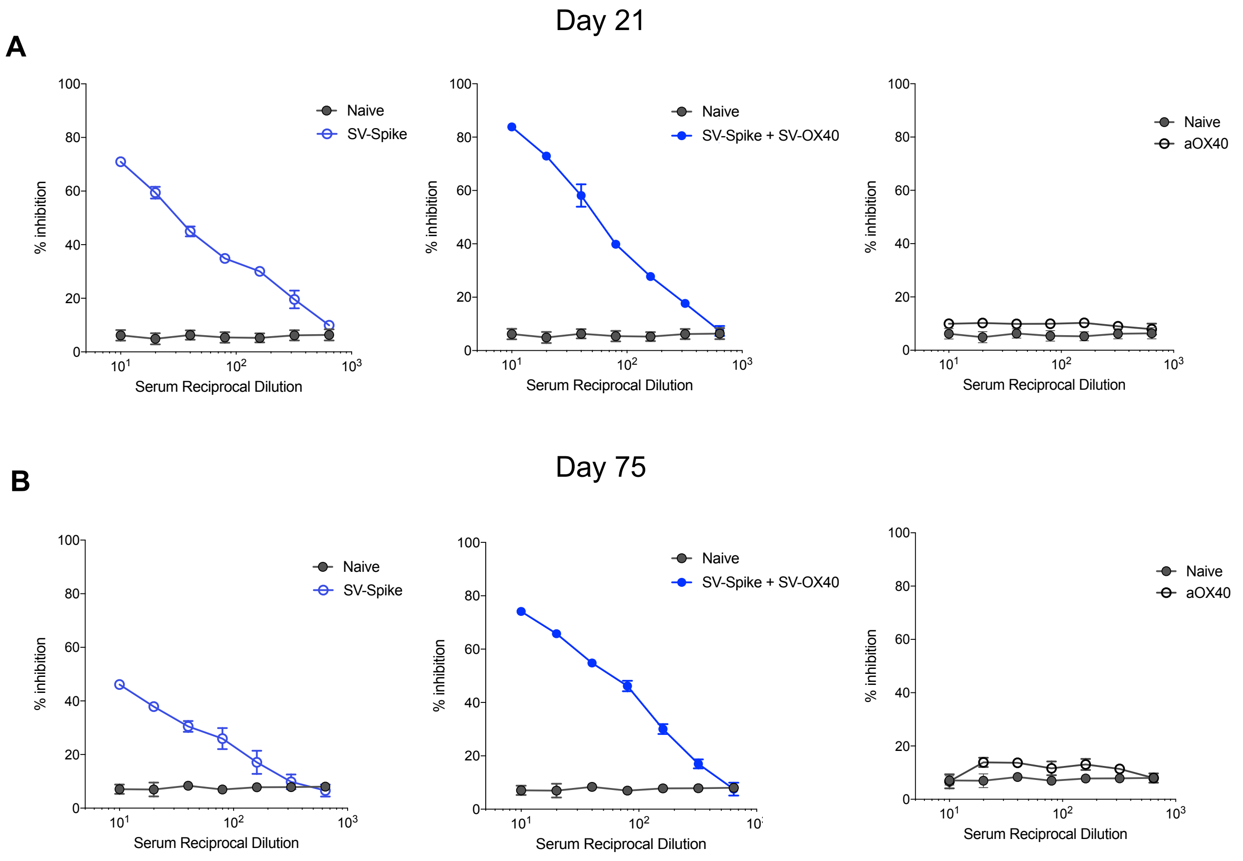


**Supplementary Figure 5.** Dose-response curves of neutralization of SARS-CoV-2 spike pseudotyped lentivirus infection by Sindbis vaccination. *Luciferase*-encoding SARS-CoV-2 pseudotyped particles were incubated with C57BL/6J mouse sera collected at **(A)** 21 and **(B)** 75 days post vaccination with Sindbis expressing SARS-CoV-2 spike (left panels), SARS-CoV-2 Spike in combination with αOX40 (middle panels) and αOX40 alone (right panels) compared to the Naïve group. The data presented are the mean of n = 5 biological replicates with n = 2 technical replicates each curve.

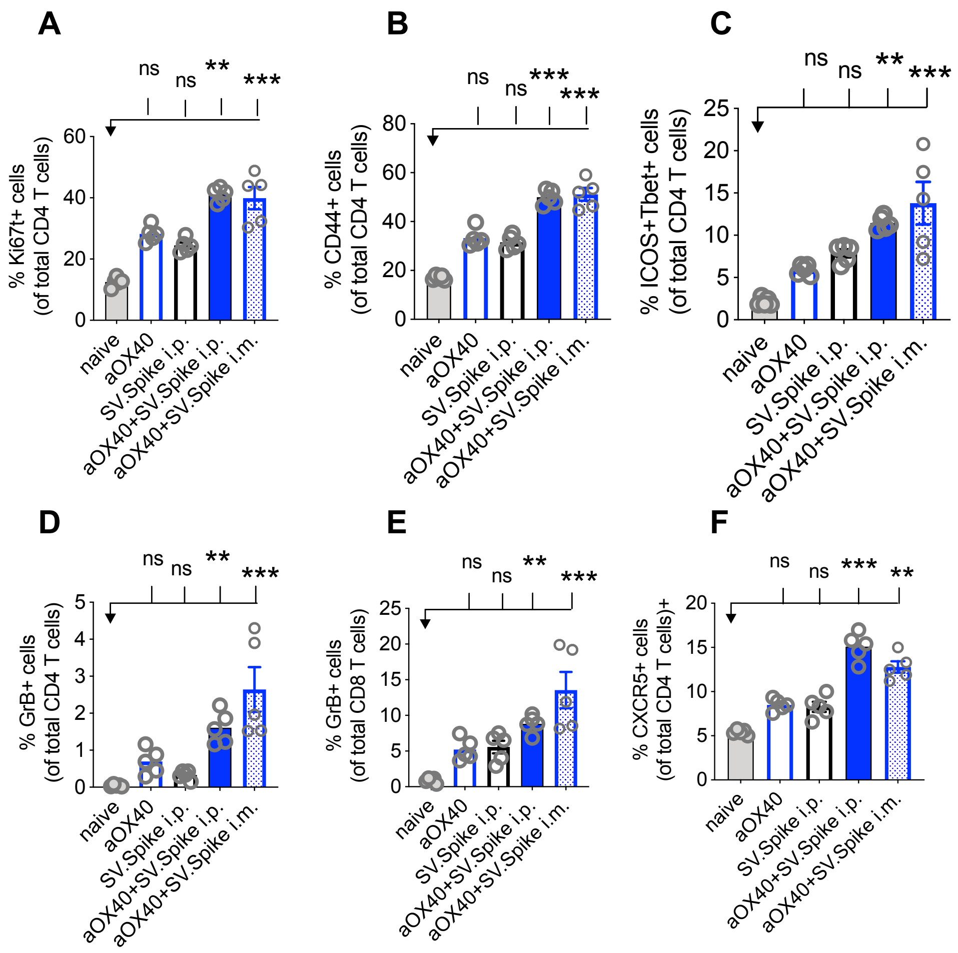


**Supplementary Figure 6.** T cell activation and differentiation. Comparison of intraperitoneal and subcutaneous immunization routes with SV.Spike in combination with αOX40. Mice were immunized with SV.Spike via the intraperitoneal or intramuscular route in combination with αOX40. Naive mice were used as control. Spleens were excised and single- cell suspensions were stained for flow cytometry analysis on day 7 after prime doses. **(A)** Proliferation of CD4 T cells indicated by Ki67+ expression. **(B)** CD4 T cell activation indicated by CD44+ expression. **(C)** Th-1 type T cell differentiation indicated by double-positive ICOS+Tbet+ expression. Cytotoxic CD4+ **(D)** and **(E)** CD8 T Cells indicated by GrB+ expression. **(F)** CXCR5+ upregulation indicates Tfh cells differentiation. Bars or symbols represent means ± SEM (n=5 mice each group). Statistical significance was determined with the Kruskal-Wallis test followed by the he Dunns' test. n.s. > 0.05, **p<0.005, ***p≤ 0.001.

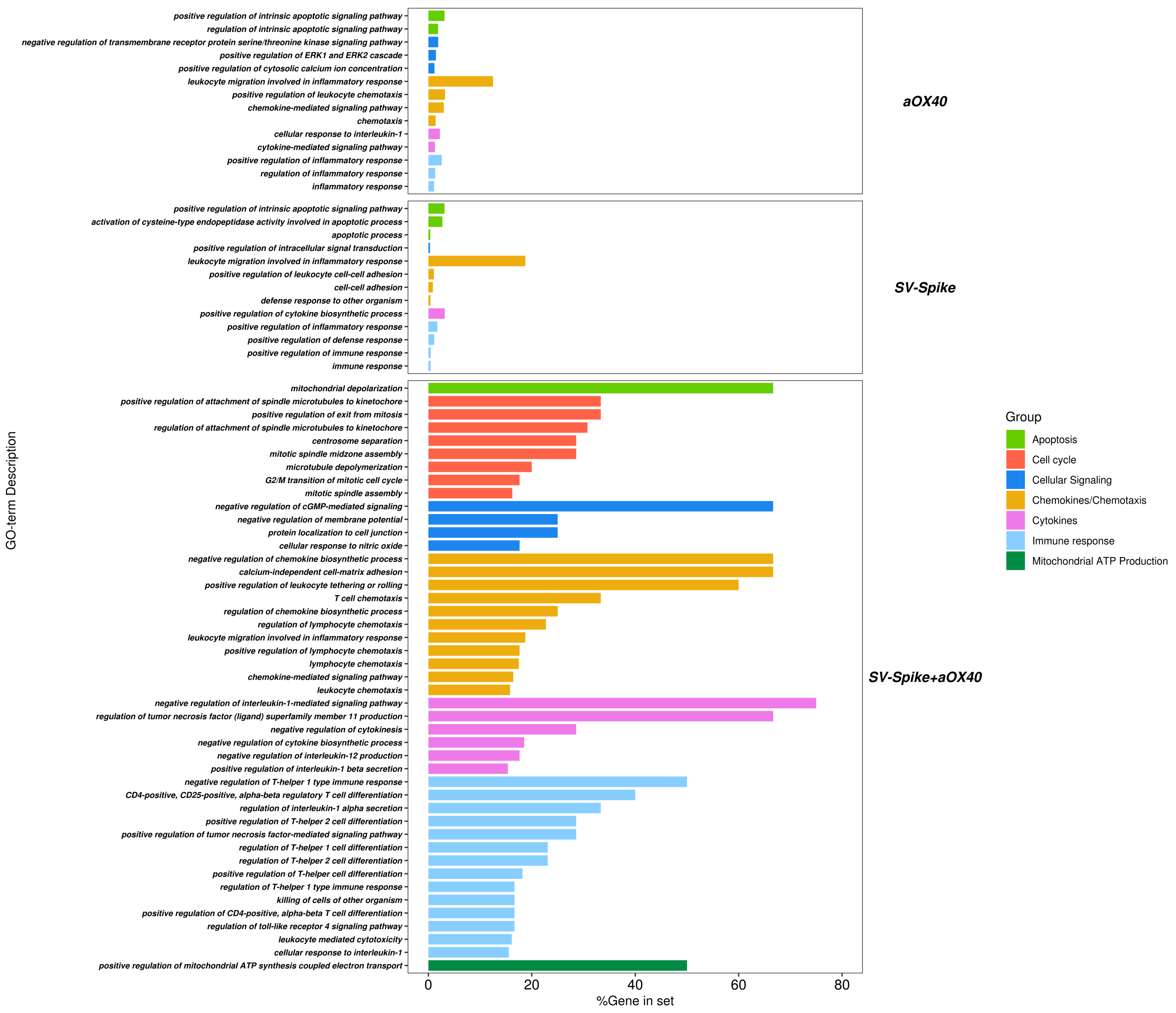


**Supplementary Figure 7.** T Cells of SV.Spike+αOX40 vaccinated mice show a unique transcriptional signature compared to single agents. Combination therapy markedly changes the transcriptome signature of T cells favoring T cell differentiation towards effector T cells shortly after prime vaccination. T cells were isolated and RNAseq was performed. Gene ontology analysis for biological processes was performed by STRING. Significantly upregulated DEGs (≥2 fold) from T cells isolated from SV.Spike and/or αOX40 treated C57BL/6J mice compared to naïve group were analyzed. Each bar represents a functional annotation (Strength ≥1). Percentage of contributing upregulated DEGs per GO term is indicated for αOX40 (top), SV.Spike (middle) and combination vaccinated group (bottom). Biological processes are further clustered for Apoptosis (light green), Cell Cycle (red), Cellular Signaling (blue), Chemokines/ Chemotaxis (orange), Cytokines (pink), Immune response (light blue) and Mitochondrial ATP Production (green).

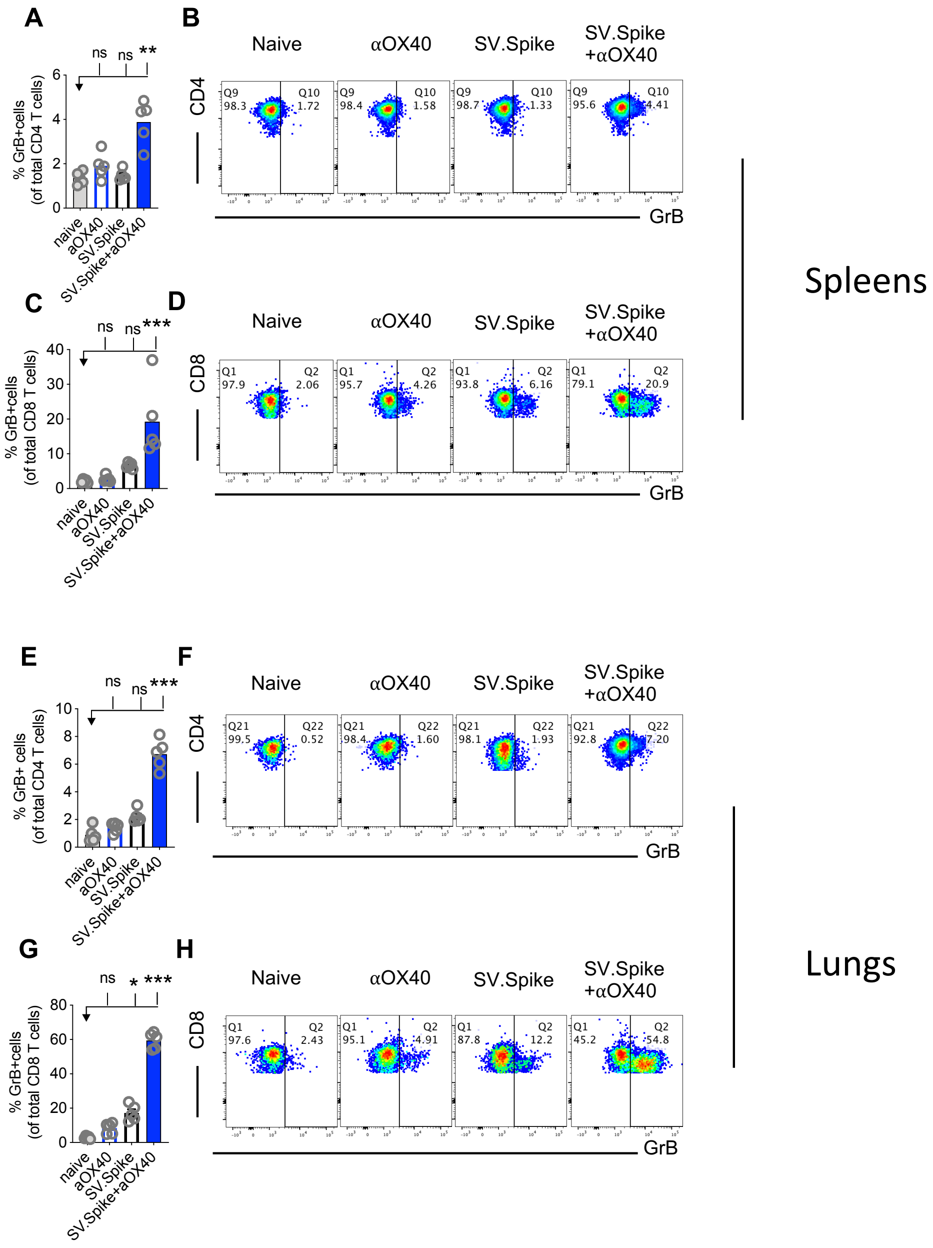


**Supplementary Figure 8**. SV.Spike in combination with αOX40 drives cytotoxic T cell differentiation. C57BL/6J mice were prime/boost immunized with SV.Spike and/or αOX40. Naive mice were used as control. Spleens **(A-D)** and lungs **(E-H)** were excised and single- cell suspensions were stained for flow cytometry analysis on day 21 after prime doses. Cytotoxic CD4+ **(A, B, E, F)** and CD8+ T cells **(C, D, G, H)** were present as indicated by granzyme B+ positive cells in spleens and lungs. (n=5 mice per group). Representative blots are shown. Bars or symbols represent means ± SEM. Statistical significance was determined with the Kruskal-Wallis test followed by the he Dunns' test. n.s. > 0.05, *p<0.05, **p<0.005, ***p≤ 0.001.

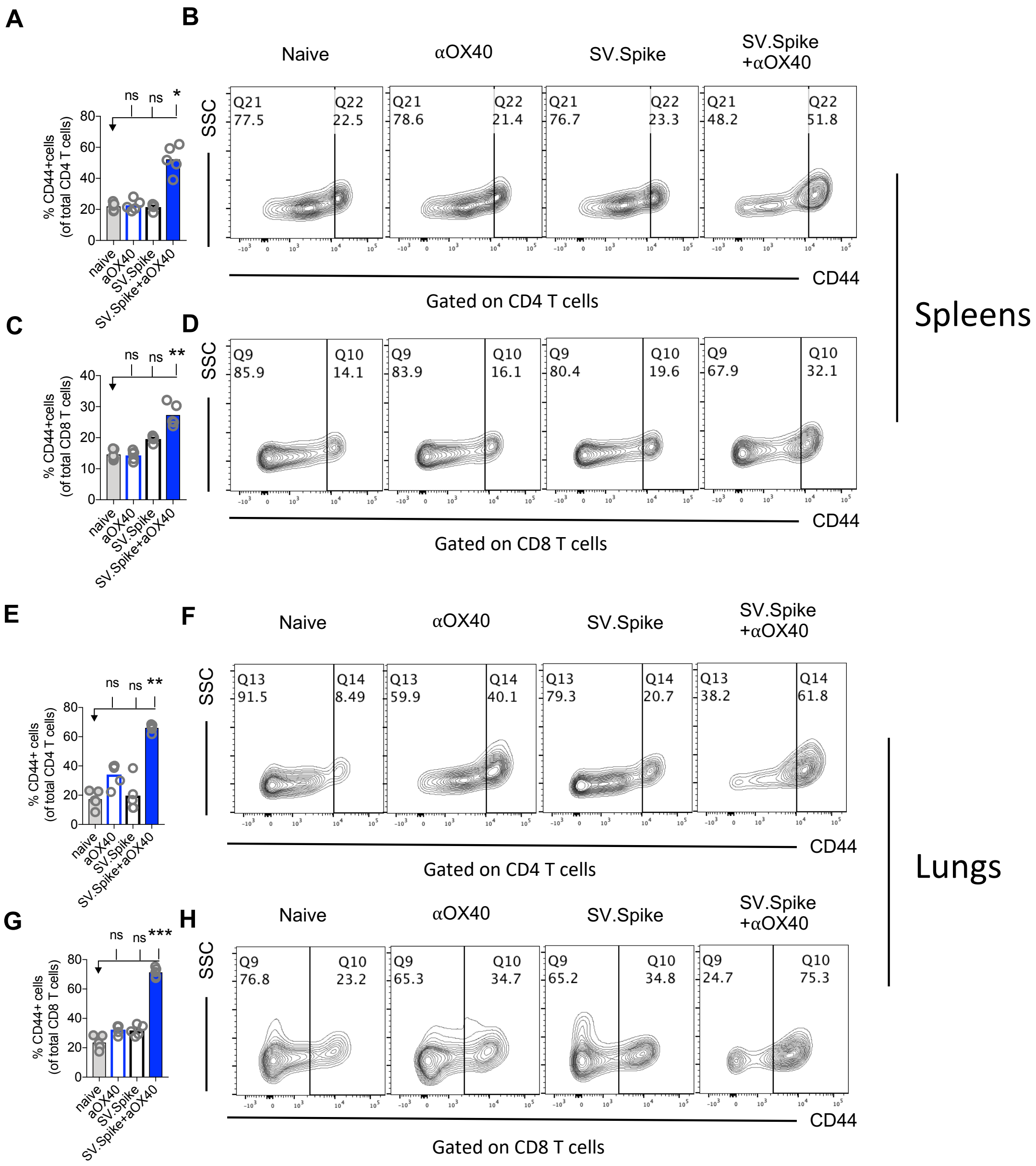


**Supplementary Figure 9.** SV.Spike in combination with αOX40 drives T cell activation. Mice were prime/boost immunized with SV.Spike and/or αOX40. Naive mice were used as control. Spleens **(A-D)** and lungs **(E-H)** were excised and single- cell suspensions were stained for flow cytometry analysis on day 21 after prime vaccine doses. Activated CD4+ **(A, B, E, F)** and CD8+ T cells **(C, D, G, H)** were present as indicated by CD44+ positive cells in spleens and lungs. Bars represent means and each symbol represent an individual mouse Statistical significance was determined with the Kruskal-Wallis test followed by the he Dunns' test. (n=5 mice per group) n.s. > 0.05, *p<0.05, **p<0.005, ***p≤ 0.001.

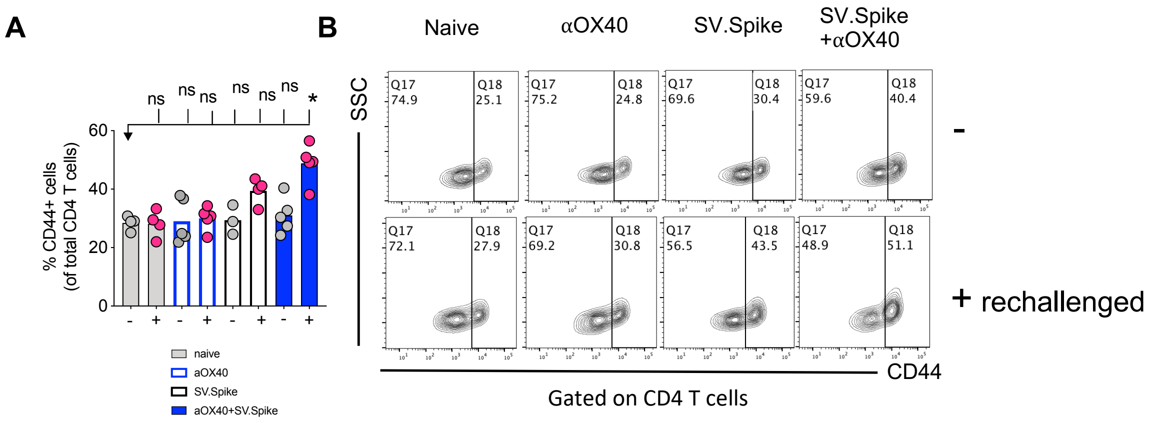


**Supplementary Figure 10.** Rechallenging immunized mice with spike antigen promotes a fast response of immune effector memory T cells. T cell activation was assessed in C57BL/6J vaccinated mice after rechallenge with Sindbis carrying SARS-CoV-2-spike. Mice were rechallenged with SARS-Cov-2 spike on day 100 after prime vaccinations. Spleens were excised on day 103 and single cell suspensions were stained for flow cytometry analysis. **(A)** CD44+ positive CD4+ T cells and representative plots **(B)** indicating T cell activation shortly after rechallenge. (n=5 mice per group, or as otherwise indicated). Bars or symbols represent means ± SEM. Statistical significance was determined with the Kruskal-Wallis test followed by the he Dunns' test. n.s. > 0.05, *p<0.05.
